# Myelin Mapping in the Human Brain Using an Empirical Extension of the Ridge Regression Theorem

**DOI:** 10.1101/2025.05.08.652955

**Authors:** Griffin S. Hampton, Ryan Neff, Zezheng Song, Mustapha Bouhrara, Radu Balan, Richard G. Spencer

## Abstract

**Purpose:** Myelin water fraction (MWF) mapping in the central nervous system is a topic of intense research activity. One framework for this requires parameter estimation from a decaying biexponential signal. However, this is often an ill-posed nonlinear problem resulting in unreliable parameter estimates. For linear least-squares (LLS) problems, the ridge regression theorem (RRT) shows that a Tikhonov regularization parameter exists that will reduce mean square error (MSE) in parameter estimates. We present and apply a nonlinear version of the RRT, *λ*-NL-RR, to MWF mapping.

**Methods:** For simulated and experimental data, we estimated parameter values with conventional nonlinear least-squares (NLLS) and compared these with values obtained from *λ*-NL-RR, with the regularization parameter value defined by generalized cross validation. We applied regularization only to signals identified as biexponential according to the Bayesian information criterion.

**Results:** Under conditions of modest SNR and closely spaced exponential time constants in which conventional biexponential analysis methods yield particularly inaccurate results, *λ*-NL-RR decreases MSE by ~10-15%.

**Conclusion:** Regularization of the NLLS parameter estimation problem for the biexponential model decreased MSE for simulated and in vivo MRI brain data. In addition, this work provides a general framework for regularization of a broad class of NLLS problems.

## 1 INTRODUCTION

### 1.1 Motivation

Myelin water fraction (MWF) mapping in the human brain is a subject of intense research activity; it requires separating the signal component arising from water molecules trapped within the myelin sheath from signal generated by less bound water [1, 2, 3]. The MWF is closely associated with white matter (WM), as compared to the largely non-myelinated grey matter (GM) [4, 5, 6, 7]. Approaches to quantifying MWF include multi spin- or gradient-echo experiments, resulting in transverse decay which can be modeled as a biexponential:

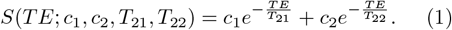

with typically *T*_21_ ∈ [10, 50] ms and *T*_22_ *>* 60 ms [5, 8, 9], *c*_1_ and *c*_2_ corresponding respectively to the myelin and non-myelin water fractions, and MWF= *c*_1_*/*(*c*_1_ + *c*_2_). *TE*, the echo time, is the independent variable for signal acquisition. For generalization to other settings, we will take the minimum value of *TE* = 0, although in MRI, *TE >* 0 due to pulse and gradient durations. This formulation is quite distinct from non-negative least squares analysis, which is not the topic of the present work [10].

Eq. 1 appears throughout mathematics and physics [10, 11, 12, 13, 14]. In spite of its apparent simplicity, parameter estimation from this model inherits the ill-posedness of the inverse Laplace transform and the Fredholm integral equation of the first kind, of which it is a special case [10, 11, 15, 16, 17, 18]. As a result, estimates can be extremely noise-sensitive, depending on the values of the time constants, component fractions, and SNR [19].

In contrast, parameter estimation from the monoexponential, single-component decay model

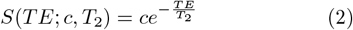

with parameter designations analogous to those for Eq. 1, is much more well-posed. This can be illustrated by the condition number of each model (Figure S1).

Regularization of linear and nonlinear inverse problems is an established technique for stabilizing parameter estimation. Although this introduces bias, it is wellknown that biased estimators can yield mean squared error (MSE) less than that of an unbiased estimator, given by the Cramér-Rao lower bound (CRLB). In general, a regularization approach for recovering, e.g., a distribution function or an image is selected based on assumed properties of the desired result. Thus, regularization can be implemented which limits solution variability, or gradients, or curvature, or total variation, or that promotes sparsity. In the present case, the desired solution is a set of unrelated parameter values, that is, delta-functions in the solution space. In fact, these delta functions are in different solution spaces: the space of possible component values with dimensions of “percent”, or “fraction”, and the space of relaxation times, with dimensions of “time”. The novelty of the present work is, therefore, that we apply regularization to the estimation of discrete parameters rather than to a discretized continuum of parameters. We rely only on the underlying mathematical properties of regularization rather than physical insight into the desired solution. In addition, we deliberately inject and adjust bias through regularization in order to design an estimator with decreased MSE. Thus, this work provides a new method for parameter estimation in biexponential modeling, and may also provide a new framework for the ubiquitous problem of least-squares parameter estimation.

### 1.2 Biexponential Analysis

Estimator performance can be quantified through bias, the statistical expectation value of the difference between parameter estimates and actual values, and variance, a measure of the variability of estimates. Variance as a metric does not require knowledge of underlying parameter values; it depends only on estimator output. However, MSE is the preferred metric for estimator quality:

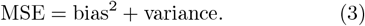

The CRLB equals MSE only for unbiased estimators, limiting its utility for nonlinear estimation except in the limit of high signal-to-noise (SNR).

### 1.3 Least-Squares Analysis for

#### Parameter Estimation

Nonlinear least-squares (NLLS) parameter estimation for 1-dimensional signals is defined by:

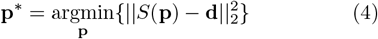

where *S*(**p**) is a possibly nonlinear signal model with parameters **p, d** is observed data, and **p**^*^ is the optimal parameter set. The bracketed objective function is called the residual sum of squares (RSS).

One goal of estimation methodology is to decrease the sensitivity of **p** to noise in **d**. A standard method is Tikhonov regularization (TikReg) [10, 20] which re-casts Eq. 4 as:

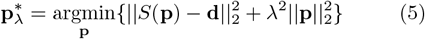

where *λ* is the regularization parameter. The difference between the objective functions of Eq. 5 and Eq. 4 introduces bias into estimates of **p**, but also reduces variance so that MSE can be reduced. There are innumerable variations of TikReg, with penalties constructed based on prior knowledge of relationships between the elements of **p**, which necessarily have the same units and dimensions. Thus, TikReg is not well-studied in the framework of Eq. 5 when **p** is a set of discrete parameters with no obvious physical relationship, and which may be of different physical dimensions. In our case, for example, there is no a priori reason for *c*_1_ *≈ c*_2_ or *T*_21_ *≈ T*_22_, or for the norm of the vector of these parameters to have small total variation or magnitude.

### 1.4 The Ridge Regression Theorem

Linear least-squares (LLS) parameter estimation is unbiased but may still be ill-posed. It is well-known that use of biased estimators, like ridge regression, can reduce MSE [21, 22, 23, 24, 25, 26, 27]. This is highlighted by the ridge regression theorem (RRT) from the statistics literature, which states that for the LLS problem of Eq. 5, there always exists a *λ >* 0 that reduces MSE [28]. Lacking a general theory for the nonlinear case, we empirically address the question of whether MSE can be reduced through bias injection via TikReg.

Selecting the optimal *λ* is a central problem in TikReg [24, 29, 30, 31]. Here, we use generalized cross validation (GCV), a popular, statistically validated method that does not require an estimate of noise amplitude [32, 33, 34, 35, 36].

### 1.5 Estimator Metrics and the Biased

#### CRLB

A common metric for estimator effectiveness is the CRLB, the minimum variance, and hence minimum MSE, achievable by an unbiased estimator [23, 26]. CRLB analysis is often used to guide experimental design even for nonlinear, and therefore biased, estimation [26, 37, 38, 39, 40, 41, 42, 43]. However, a CRLB for biased estimation, the biased CRLB (bCRLB), can also be calculated and defines the minimum variance for a biased estimator [40, 44]. It is derived from a bias gradient matrix that, like the conventional CRLB, is specific to the estimator, model parameters, and noise [25, 40, 45, 46]. Importantly for our purposes, the bCRLB can also incorporate regularization, unlike the conventional CRLB. Therefore, we can establish three metrics for comparison with MSE: the conventional CRLB and bCRLB for the nonregularized estimator, and the bCRLB for the regularized estimator. Quantification of bias requires knowledge of underlying parameter values which are unknown in applications, but are available in simulations.

### 1.6 Model Selection for Regularization of Brain Data

Although a *λ*-selection technique such as GCV would ideally return a value of *λ ≈* 0 when estimator variance is already low, all *λ* selection methods are imperfect. GCV often exhibits a very flat curve of GCV score vs. *λ*, so that a well-posed problem exhibiting low noise sensitivity may be over-regularized. GM, representing minimally- or non-myelinated cellular tissue, is expected to exhibit monoexponential decay, from which parameter extraction is well-posed. In contrast, the myelinated WM is expected to exhibit biexponential decay. Therefore, we implemented a pre-processing step in which the decay curve for each voxel was assigned as either bi- or monoexponential using the Bayesian information criterion (BIC). Since the BIC is parsimonious in defining model complexity, this reserves regularization for curves that are strongly biexponential [47, 48].

## 2. METHODS AND DATA

### 2.1 Simulated Data

#### 2.1.1 Weighted NLLS with Regularization

We introduced a diagonal weighting matrix *W* = diag[1, 1, 1*/*100, 1*/*100] in Eq. 5 to ensure roughly equal contributions from the four parameters to be estimated [8, 9]:

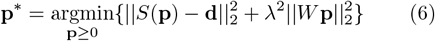

We refer to this nonlinear parameter estimation problem, based on the RRT, as *λ*-NL-RR.

#### 2.1.2 Generating Simulated Signals

Signals were generated according to Eq. 1 for parameter values in the expected range for brain (Table 1), with 128 echo times (*TE*) from 0 to 635 ms in steps of 5 ms. This ensured full sampling of the signal decay, given typical *T*_2_ values for GM of ~80 ms, and ~10 to 50 ms respectively for the WM myelin-associated water and ~100 ms for relatively mobile water in WM [8, 9]. We used *TE*_0_ = 0 for general applicability; the modification to a small but non-zero *TE*_0_ does not impact our analysis or results in any substantial way (not shown).

**TABLE 1.** Parameter combinations resulting in 225 noiseless signal curves, from which parameters were derived with the indicated SNR values.

| Parameter | Values |
| --- | --- |
| $(c_1, c_2)$ | (0.0, 1.0), (0.05, 0.95), (0.2, 0.8),<br>(0.4, 0.6), (0.5, 0.5), (0.6, 0.4),<br>(0.8, 0.2), (0.95, 0.05), (1.0, 0.0) |
| $T_{21}$ | 10, 20, 30, 40, 50 |
| $T_{22}$ | 70, 90, 110, 130, 150 |
| SNR | 20, 50, 100, 500 |

Signals corresponding to 225 different parameter combinations (Table 1) were generated, each corrupted by 500 independent realizations of zero-mean Gaussian noise to achieve a specified SNR, defined as the signal amplitude at *TE*_0_, equal to 1 in this case, divided by *σ*: 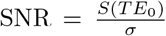. For given parameters, the same random noise matrix was used across SNR values but scaled to achieve the desired SNR. This ensured that results as a function of SNR were indicative of estimator performance rather than differences in noise realizations.

#### 2.1.3 Simulation Parameter Estimation

Estimates were performed using the trust-region algorithm within the curve fit routine from the scipy.optimize package in Python 3, using the stacked form of Eq. 6 [29]. The parameter search space was restricted to 0 *< c*_1_, *c*_2_ *<* 2, 0 *< T*_21_ *<* 100, and 0 *< T*_22_ *<* 300, based on previous estimates of MWF [5, 8, 49]. To mitigate the effect of nonconvexity of the biexponential objective function, ten random combinations of starting values, denoted with subscript 0, were used for each NLLS estimate. *c*_1,0_ was sampled from U(0,1), *T*_21,0_ from U(0,100), *T*_22,0_ from U(*T*_21,0_,300), where U denotes the uniform distribution, and *c*_2,0_ was set to 1 − *c*_1,0_. Of the ten estimates, the one that minimized the objective function of Eq. 6 was selected as the final result. By convention, the larger of the two estimated decay constant values was taken as *T*_22_, with its corresponding *c*_2_ [49]. The requirement that *c*_1_ + *c*_2_ = 1 was enforced to ensure correct interpretation as fractions of the two assumed components. This normalization was omitted when estimates were compared to CRLB analysis, which was based on non-normalized values.

#### 2.1.4 Cramér-Rao Lower Bound

We calculated the CRLB matrix for specified combinations of parameters **p** [19, 50]:

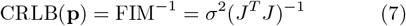

where FIM denotes the Fisher information matrix, *J* is the Jacobian of the biexponential model, 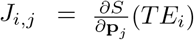, and *σ* is the standard deviation of the noise. For the case of additive white Gaussian noise (AWGN) considered here, the CRLB for each parameter is the diagonal element of the CRLB matrix corresponding to the ordering of **p**.

We also computed the bCRLB [25, 44], which requires calculation of the bias gradient:

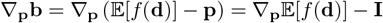

where the first expression inside the derivative is the expectation value of the bias for the estimator *f* (**d**). The measured data is given by **d** = *S* + *ν*, with *S* from (1) and *ν*defining the AWGN. The data and probability density functions, *p*_**d**_(**d**) and *p*_*ν*_ (*ν*), are related via *p*_**d**_(**d**) = *p*_*ν*_ (**d** − *S*).

The required Jacobian matrix *R* = ∇_**p**𝔼_ [*f* (**d**)] was computed numerically using stochastic Monte Carlo integration of

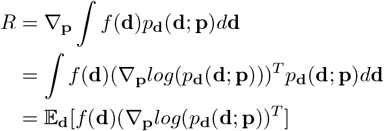

where the expectation 𝔼_**d**_ is with respect to the measured data **d**. In deriving the first step, note that *f* (**d**), the estimator, acts on given data **d** and returns a four-vector of parameters; in this sense, while **d** depends on **p**, the estimator *f* (**d**) depends only on **d** and not on **p**.

The gradient of the log-likelihood can be computed analytically for AWGN:

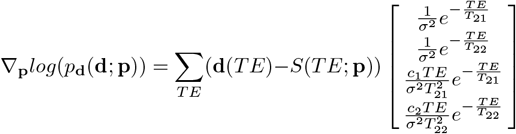

For the nonregularized NLLS estimator, we simulated and averaged over 10^7^ noise realizations. For the GCV estimator, we simulated and averaged over 10^4^ noise realizations due to its higher computational cost.

We validated these estimates using a second method for computing the Jacobian. For a given **p**_0_ we generated 256 random 4-vectors in the neighborhood **p**_0_ ± [0.05, 0.05, 2.5, 2.5], and for each we simulated and averaged over 10^5^ noise realizations. Then the *R* matrix is obtained as the least-squares solution over these 256 estimates in the model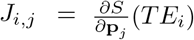. We estimated *R* for two sets of parameters and two levels of SNR. The bCRLB for each parameter was computed as the diagonal element from the matrix determined from Eq. 6 of [44]:

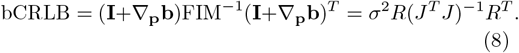

This also follows directly from Eq. 3.30 of [50], with the function *g* there taken as *g*(**p**) = **p** + **b**(**p**), where **b**(**p**) is bias. We denote bCRLB values by bCRLB_Nonreg_ when bias is due solely to the presence of noise in the nonlinear estimation, and bCRLB_Reg_ when bias is further introduced by *λ*-NL-RR.

#### 2.1.5 Oracle Method of Regularization Parameter *λ*

We implemented an “oracle” method of *λ* selection as a benchmark for optimal performance of *λ*-NL-RR. We calculated the average MSE across 500 noise realizations for 51 logarithmically-spaced *λ* values from 10^−7^ to 10, along with the zero value. The oracle lambda is the value for which MSE is the minimum. Plots of MSE versus *λ* indicate whether values exist for which MSE is decreased below that of nonregularized NLLS analysis. This visualization confirms the validity of the RRT in this setting, so that it is meaningful to seek a regularization scheme to decrease MSE, although it is, of course, not helpful in *λ* selection.

#### 2.1.6 Generalized Cross Validation *λ* Selection

We implemented a GCV method for nonlinear inverse problems [36]; GCV identifies the *λ* which minimizes:

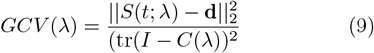

Where

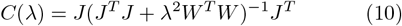

given a signal **d** with number of echo times *n*, a regularized model *S*(*t*; *λ*), its Jacobian *J*, and the weight matrix *W* as defined above. *GCV* (*λ*) is noise-dependent, so that a different *λ* is selected for each noise realization. The same array of *λ*s as for the oracle calculation was sampled for GCV *λ* selection. For *λ* = 0, the *C*(*λ*) matrix reduces to *J* (*J*^*T*^ *J*)^−1^*J*^*T*^ and is sometimes singular. In such cases, the denominator in the *GCV* (*λ*) expression was set to the difference between the number of data points and the rank of the Jacobian. Thus, for log_10_ *λ <* −7, we replaced the denominator of Eq. 9 with (*n* − rank*J*)^2^. This comes about by noting that when *λ* = 0, *C*(*λ*) becomes an orthogonal projection matrix, that is, idempotent and symmetric; it is well-known that the trace of such a matrix equals its rank. We added this exception to our GCV method to ensure that *λ* = 0 maintains the expected GCV value when *J* is singular, namely, when *T*_21_ *≈ T*_22_. Note that in this case, the rank of *J* is further reduced from 3 to 2 if *c*_1_ = *c*_2_. In such cases, although *J*^*T*^ *J* is square, it is singular and hence noninvertible. The indicated inverse is then replaced with the Moore-Penrose pseudoinverse.

#### 2.1.7 Evaluation Metric for Simulated Data

For each parameter combination, SNR level, and regularization parameter, we determined estimates 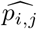 for the *j*^th^ parameter of **p** over 500 noise realizations labeled by *i*. The average estimated parameter value and bias were calculated respectively as 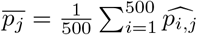 and

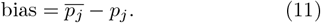

Variances were calculated as:

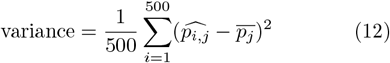

We then used Eq. 3 to calculate MSE.

The effectiveness of λ-NL-RR was defined by

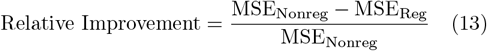

where the MSE for the conventional (nonregularized) and regularized NLLS analyses are denoted MSE_Nonreg_ and MSE_Reg_, respectively. We replaced MSE with bias and variance in Eq. 13 to calculate analogous metrics for these quantities.

### 2.2 Experimental Data

#### 2.2.1 Brain Data

3D gradient and spin-echo (GRASE) brain images were obtained from a healthy 41-year-old male, m41, and a healthy 79-year-old male, m79, using a 3T Philips MRI system (Achieva, Best, the Netherlands), with a quadrature body coil for transmission and an eight-channel phased-array head coil for reception. Parameters included 32 data acquisition times with initial value and spacing of 11.3 ms, TR = 1000 ms, echo planar imaging acceleration factor of 3, field of view 278 mm x 200 mm x 30 mm, acquisition matrix 185 x 133 x 10, and acquisition voxel = 1.5 mm x 1.5 mm x 3 mm reconstructed to = 1 mm x 1 mm x 3 mm. Scan time was approximately 10 minutes. Examinations were in compliance with the standards of our Institutional Review Board. We evaluated a central mid-volume axial slice for each subject. Additional slices are included in the SI. After skull-stripping, voxels containing primarily CSF were excluded by thresholding.

Reference images were created using the NESMA non-local denoising filter [49]. Zero-mean Gaussian noise was then added to achieve desired SNR. A constant parameter was added to Eqs. 1 and 2 to partly account for the Rician noise floor. We defined reference image noise standard deviation *σ*_Ref_ as the standard deviation of signal values at maximum *TE* from a region of interest (ROI) of approximately uniform intensity within the left-sid WM. The ROI was 18 x 55 voxels for m41 and 13 x 45 voxels for m79. Mean initial signal amplitude 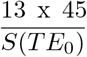 was defined over the same region. After adding noise, the initial decay amplitude for each voxel was normalized to one. For m41, the reference image exhibited an SNR of 159, and noise was added to obtain data with SNR values of 100 and 75. SNR for the reference image for m79 was 102, with noise added to obtain data with SNR of 75.

#### 2.2.2 BIC Filter Application

Each brain voxel decay signal was classified as biexponential or monoexponential using the BIC [51]:

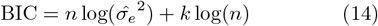

where *n* is the number of data points, *k* is the number of model parameters (3 or 5 for mono- or biexponential, including noise offset term), and 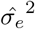 is the maximum likelihood estimate of variance:

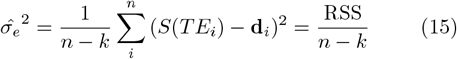

. Then:

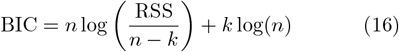

and the model with the smaller BIC is selected.

#### 2.2.3 Experimental Parameter Estimation

GCV was implemented with 101 *λ* values on a logarithmic scale from 10^−5^ to 10, along with the zero value, for regularization of each voxel identified as exhibiting biexponential decay according to the BIC.

As noted, decay signals were normalized to one before fitting. For monoexponential fits, the parameter search space was restricted to 0 *< c <* 1.5 and 0 *< T*_2_ *<* 300 with initial parameters *c*_0_ = 1 and *T*_2_ = 20. For biexponential signals, the space was restricted to 0 *< c*_1_ *<* 0.75, 0 *< c*_2_ *<* 2, 0 *< T*_21_ *<* 80, and 0 *< T*_22_ *<* 300 with initial parameters *c*_1,0_ = 0.2, *c*_2,0_ = 0.8, *T*_21,0_ = 20 and *T*_22,0_ = 80, based on typical MWF mapping analyses [5, 8]. For both models, the offset parameter was constrained to positivity. After parameter estimation, *c*_1_ and *c*_2_ were normalized to sum to one. *T*_21_ and *T*_22_, along with the corresponding *c*_1_ and *c*_2_ values, were ordered according to the convention *T*_21_ *< T*_22_.

We used parameter values obtained from the NESMA-filtered images using conventional NLLS as gold standards. Metrics, including MSE, obtained across noise realizations were calculated relative to these standards.

#### 2.2.4 Evaluation Metrics for Brain Data

We compared results obtained using *λ*-NL-RR with those from conventional NLLS for the estimation of *c*_1_, representing the MWF. Improvement in RMSE for each voxel *i* across 20 noise realizations, denoted ΔRMSE_all,*i*_, was defined as:

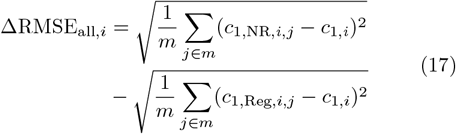

where *m* is the number of noise realizations, labeled by *j*, for which a given pixel was found to be biexponential according to the BIC. *m* can vary across noise realizations and so may differ across voxels.

## 3 RESULTS

### 3.1 Simulation Analysis

#### 3.1.1 Effect of Regularization

Figure 1 demonstrates that the basic properties of *λ*-NL-RR are consistent with the bias-variance behavior expected from regularization [40]. For the ill-posed example shown, *c*_1_ and *c*_2_ were fixed, and two-parameter estimation of *T*_21_ and *T*_22_ was performed. Contours, with minima set to zero to facilitate comparison, show level sets of the full objective function (native plus regularizer) in Eq. 6, as generated by the (*T*_21_, *T*_22_) pairs without noise. The red dot shows the underlying values of *T*_21_ and *T*_22_. Noise was added to each signal to achieve SNR=100, with the green dots showing the (*T*_21_, *T*_22_) pairs that minimize the objective function over 1000 noise realizations. Moving from Panel **(A)** (nonregularized) to Panels (*λ*=0.2) and **(C)** (*λ*=0.4) shows the decrease in variance and increase in bias with increased regularization. A well-posed case is shown in Figure S2.

**FIGURE 1.**
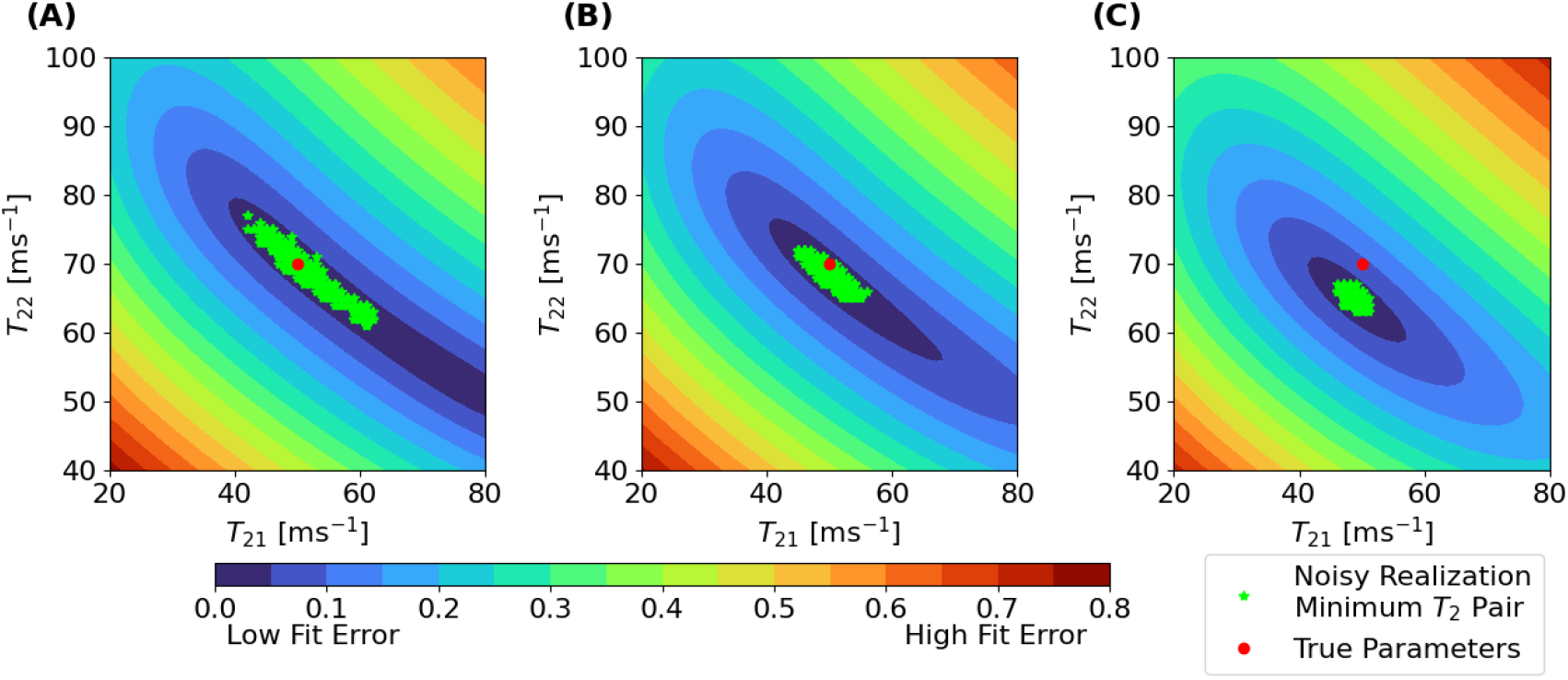
NLLS Objective Function Surface for a Nonregularized and Regularized Ill-Posed Problem. Contours show equal-value surfaces of the regularized objective function in Eq. 6, with model parameters [*c*_1_, *c*_2_, *T*_21_, *T*_22_] = [0.4, 0.6, 50, 70]. The red dot indicates these underlying values of *T*_21_ and *T*_22_. For the underlying values of *c*_1_ and *c*_2_, and with *T*_21_ and *T*_22_ varying as indicated in the figure, a decay curve was generated to which noise was then added to achieve SNR = 100. Then for each of 1000 noise realizations, a green dot is generated indicating the minimum value of the regularized objective function of Eq. 6, indicating the solution to the regularized least-squares problem. Panel **(A)**: *λ* = 0 (nonregularized). The centroid of the green dots is slightly displaced from the red dot, indicating the underlying bias of this nonlinear estimation problem even in the absence of regularization. Panel **(B)**: *λ* = 0.2, with clear displacement of the centroid of solutions from the correct value, but with decreased variance of the set of solutions to the noisy estimation problem. Panel **(C)**: *λ* = 0.4, illustrating still greater bias and a further reduction of variance.

#### 3.1.2 Empirical Demonstration of the Ridge Regression Theorem

Figure 2 compares metrics for an ill-posed problem with relatively low SNR. There is a range of *λ* for which MSE(*λ*) *<* MSE_Nonreg_, with the minimum, MSE_Oracle_, achieved by the oracle *λ. λ*-NL-RR is useful only if a *λ* can be found such that MSE_Reg_ *<* MSE_Nonreg_. The three CRLB bounds on variance are also shown. Consistent with Eq. 3, bCRLB_Reg_ *<* MSE_Reg_ and bCRLB_Nonreg_ *<* MSE_Nonreg_. We also see that the CRLB *>* bCRLB, as well as larger than the three calculated MSE values, indicating that CRLB may not be an ideal target for experimental design for nonlinear models.

#### 3.1.3 Effect of Parameter Values and SNR

We expect *λ*-NL-RR to show greater effectiveness for more ill-posed problems. For well-posed problems, increased bias can outweigh any further marginal decrease in variance. This is shown in Figure 3, showing estimation of *c*_1_ as a function of *T*_21_, *T*_22_ and SNR. The effectiveness of *λ*-NL-RR is shown by the difference between MSE_Nonreg_ and MSE_Reg_. In the problematic regime of *T*_21_ ~ *T*_22_ and modest SNR (Panel **(A)**), MSE is substantially reduced by regularization. In contrast, for widely-separated *T*_21_ and *T*_22_ and high SNR (Panel **(D)**), all displayed bounds on MSE are small and approximately equal, and there is no regime where regularization improves estimation. Panels **(B)** and **(C)** show results respectively for parameters defining an ill-posed estimation problem for high SNR, and well-posed estimation with low SNR. In **(B)**, regularization improves estimation, in spite of the already-excellent performance of NLLS. In contrast, **(C)** shows no improvement. Thus, ill-posedness defines the regime of efficacy of *λ*-NL-RR to a greater extent than SNR.

#### 3.1.4 Trends Across Parameter Combinations

Each panel of Figure 4 shows results for *c*_1_ estimation for a given *c*_1_ across a range of *T*_21_ and *T*_22_. The bold values show relative MSE improvement (Eq. 13), with a maximum possible value of one and a negative value indicating an increase in MSE. Further, deeper orange indicates greater improvement (decrease) in MSE. The middle and lower values in each cell are the improvement in square bias and improvement in variance, respectively, using expressions analagous to Eq. 13. These relative metrics, individually averaged over noise realizations, are not, mathematically, the components of relative MSE.

The improvement through *λ*-NL-RR is clear viewing each panel from upper right to the lower left, corresponding to decreasing separation between *T*_21_ and *T*_22_. In the most well-posed region (upper right, Panel **(C)**), injected bias dominates reduction of already-small variance, leading to increased MSE. Similarly, improvement increases moving downward in columns and leftward in rows. Estimation is more difficult for smaller values of *c*_1_(Panel **(A)**), corresponding to the regime of greatest improvement through *λ*-NL-RR. Panel **(A)** also shows reduction in both bias and variance with regularization, while Panel **(C)** shows negative values corresponding to increased variance. This arises as follows. TikReg with a particular value of *λ* will always decrease variance for a particular estimation problem; however, the heatmap values are averages across 500 separately-regularized noise realizations. There is no guarantee that the variance across these will decrease as compared to the variance resulting from nonregularized analysis, particularly in cases when the latter is already low. In addition, the low variance in the denominator of this relative variance metric leads to small changes appearing as large values.

Figures S3-6 of the SI show similar patterns. As SNR increases, the opportunity for regularization to improve MSE decreases, and there are fewer (*T*_21_, *T*_22_) pairs that show improvement.

### 3.2 Brain Analysis

#### 3.2.1 Brain Region Exponentiality

As described above, the BIC assigns pixels to mono- or biexponential decay for each noise realization. GM pixels were monoexponential for virtually all noise realizations, while WM pixels were consistently biexponential. Other pixels exhibited a more mixed character. See Figure S7.

#### 3.2.2 Brain MWF Estimation

We performed *c*_1_ estimation across the brain slice for 20 noise realizations for m41 for SNR values of 100 and 75 and for m79 for an SNR of 75. These MWF estimates are shown for single noise realizations in Figure 5. The range of MWF estimates is primarily within the appropriate physiological range of 0 to 0.4 [5, 8]: average standard reference MWF values of the voxels less than 0.4 are 0.19 and 0.18 for m41 and m79. Qualitatively, regularization (Panels **(G-I)**) results in less fine-scale variability across the MWF map (Panels **(D-F)**). We visualize these differences in Figure S8 and quantify them further below. Other axial slices of the brain are characterized in Figures S9-12.

#### 3.2.3 Reduction in MSE of Brain MWF Mapping

We evaluated the efficacy of *λ*-NL-RR in the estimation of MWF by computing ΔRMSE_all,*i*_ (Eq. 17); see Figure 6. The dominant blue color corresponds to improvement of *c*_1_ estimation with *λ*-NL-RR, while red corresponds to worsening. In Panel **(A)**, the average ΔRMSE for m41 at an SNR of 100 is 0.025, corresponding to a 13% improvement in MSE. Comparable results for a single noise realization are shown in Figure S13, showing ~18% improvement in MSE. For the lower SNR value of SNR of 75, the average ΔRMSE for m41 was 0.024 in Panel **(B)** and 0.026 for m79 in Panel **(C)**, indicating an improvement of 13% for m41 and 14% for m79 for SNR of 75. The influence of regularization on standard deviation and absolute bias is shown in Figures S14 and S15. Maps of the change in MSE for additional slices can be seen in Figures S16 and S17. We summarize all metrics for improvement in MSE in Table S1.

## 4 DISCUSSION

We have shown substantial improvement in myelin mapping through TikReg of the corresponding ill-posed NLLS problem for estimation of discrete unrelated parameters. We demonstrated that GCV is reliably able to define a *λ* such that MSE(*λ*) *<* MSE_Nonreg_, rendering this a practical method. These results were compared to CRLB and bCRLB analysis, indicating their limitations for experimental design. Simulations over a wide variety of parameter combinations provide results applicable to MWF mapping and illustrate *λ*-NL-RR for different degrees of ill-posedness and SNR ranges (Figure 4). We are unaware of other analyses of the validity of the RRT for similar NLLS problems. We then applied *λ*-NL-RR to MWF mapping with multi-echo data (Figures 5 and 6) [5, 8, 52]. Qualitative and quantitative metrics indicate the success of the method in reducing error.

A novel component of our analysis was the use of the BIC to restrict *λ*-NL-RR to pixels more likely to benefit. In effect, this provides a distinction between ill- and well-posed regimes. In the CNS, these correspond roughly to myelinated and non-myelinated tissue, respectively [53]. We suggest that in similar problems with nested models, the BIC or a similar criterion may be applied as part of the analysis protocol.

Regularization is a longstanding method for stabilizing parameter estimation for ill-posed problems [24], including in biomedical research and imaging. In general, the parameter set to be recovered represents a single type of quantity, for example, in MWF analysis using non-negative least squares, where a distribution of *T*_2_ values is estimated. In these settings, there is a physical rationale behind the various TikReg schemes, incorporating various penalty norms, roughening matrices, and other types of prior information.

In contrast, we are unaware of previous use of regularization in low dimensional NLLS with incommensurate variables. There is no physical basis for such regularization: the parameters are discrete, have magnitudes that are not necessarily related or even of the same physical dimensions, and exhibit no natural ordering to define a smoothness or variability constraint. Nevertheless, the mathematical basis for stabilization with TikReg remains valid [54], permitting substantial improvement in MSE. This approach may have widespread applicability to other ill-posed least-squares and related parameter estimation analyses.

The condition number (CN) of the design matrix in LLS analysis characterizes the linear dependence among its columns, with a larger CN indicating greater linear dependence. CN is also the factor by which measurement error is amplified into errors in parameter estimation.

In contrast, nonlinear estimation problems are defined in terms of a vector-valued function of multiple variables and conditioning [10, 19] can be defined by the Jacobian of the underlying model function. In our analysis, the estimation problem exhibits worsening conditioning as *T*_21_ and *T*_22_ become more similar [19]. We expect regularization to be of particular utility in this regime, as documented in Figure 4. Further, since poor conditioning relates directly to instability with respect to noise, regularization should be of greater benefit for worsening SNR, as indeed seen in Figures S3-6.

The reduction in variance from TikReg for LLS estimation can be calculated from the closed-form expression for the covariance matrix of parameter estimates [10] as a function of *λ*. Reduction in variance for NLLS is less obvious. The solution of Eq. 6 for large *λ* is clearly a vector of zeros, with zero variance. While this limit is of no practical value, it indicated how increasing *λ* can reduce variance. The contribution of decreased variance to the decrease in MSE resulting from regularization is detailed in Figure S18.

Our results suggest that the bCRLB may be a more suitable benchmark for experimental design than the CRLB [39, 42, 43] for nonlinear problems, especially with regularization. The difference between these in typical cases remains to be systematically studied. More importantly, we find that regularization can substantially reduce MSE. Unlike variance, MSE cannot be experimentally determined, since it depends on knowledge of underlying parameter values. Nevertheless, simulations can be based on presumed parameter bounds. In fact, the CRLB is also not well-defined by experimental data, since the derivatives are evaluated at estimated but unknown parameter values. More generally, optimizing experimental design around the CRLB, or even the bCRLB, may miss substantial improvements resulting from design based on MSE and incorporating regularization.

Estimator benchmarks are compared in Figs. 2 and 3. The interpretation of the CRLB for a biased estimator is unclear, and there does not appear to be a provable relationship between it and bCRLB_Nonreg_ or bCRLB_Reg_. The relationship between these two latter quantities is also unclear. In the limit of large *λ*, bCRLB_Reg_ will approach zero. However, even with oracle selection of *λ* for minimizing MSE(*λ*), there is no obvious rationale to claim that the minimum variance of the optimally regularized problem is smaller than that of the nonregularized problem. In contrast, bCRLB_Nonreg_ ≤ MSE_Nonreg_, and similarly bCRLB_Reg_ ≤ MSE_Reg_.

**FIGURE 2.**
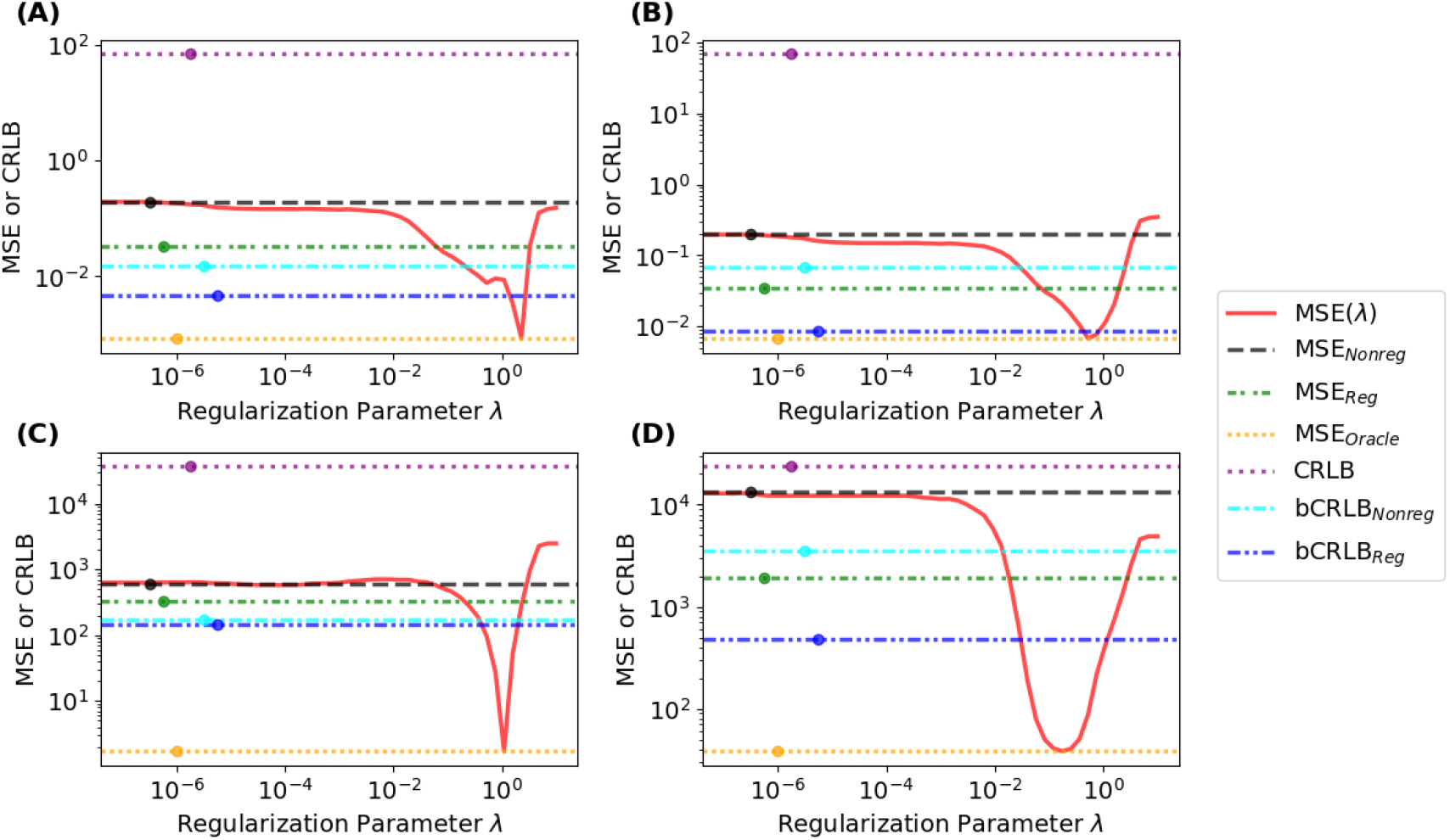
Effect of Regularization on MSE and CRLB Metrics for All Estimated Parameters. Estimator metrics averaged over 500 noise realizations for underlying parameters: [*c*_1_, *c*_2_, *T*_21_, *T*_22_] = [0.4, 0.6, 50, 70] with SNR = 20. Panel **(A)**: MSE of *c*_1_. Panel**(B)**: MSE of *c*_2_. Panel **(C)**: MSE of *T*_21_. Panel **(D)**: MSE of *T*_22_. MSE(*λ*) is shown and compared with MSE_Nonreg_, MSE_Reg_ as determined by GCV, and MSE_Oracle_, the minimum MSE achievable with *λ*-NL-RR. We also show the conventional CRLS and the biased CRLB values with and without regularization, bCRLB_Nonreg_, and bCRLB_Reg_, with GCV-defined *λ*. MSE_Oracle_ is the smallest possible MSE, but is non-realizable in practice. As expected, for small values of *λ*, MSE(*λ*) = MSE_Nonreg_. As *λ* increases, all recovered parameter values approach zero, with bias therefore defined by their ground truth values, and variance approaches zero. This accounts for the increase in MSE(*λ*) for this regime. However, for all parameters, there is a range of *λ* for which MSE(*λ*) *<* MSE_Nonreg_, indicating the success of *λ*-NL-RR. While there does not appear to be a provable relationship between the three CRLB calculations, we find that CRLB *>* bCRLB_Nonreg_ *>* bCRLB_Reg_ so that accounting for bias reduces the calculated CRLB. The indicator dots are placed on the horizontal lines for clarity due to overlapping values of the horizontal lines, with their vertical position showing ordinate values. The horizontal position of these dots is varied only for display purposes and is of no significance.

**FIGURE 3.**
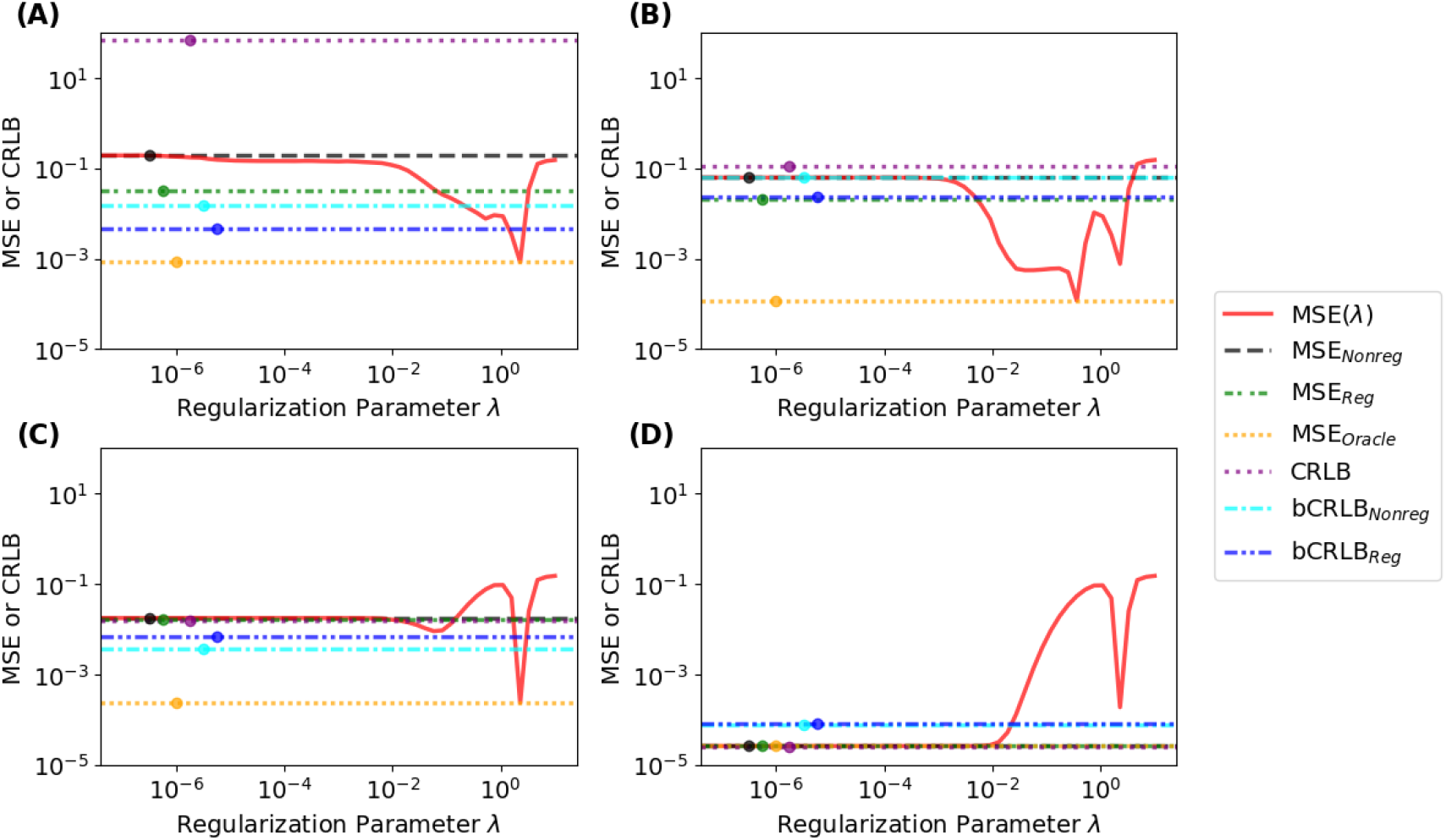
Effect of Regularization on MSE and CRLB of MWF Estimation as a Function of Ill-posedness and SNR. Estimator metrics averaged over 500 noise realizations for underlying parameters. Panel **(A)**: [*c*_1_, *c*_2_, *T*_21_, *T*_22_] = [0.4, 0.6, 50, 70] and SNR = 20. Panel**(B)**: [0.4, 0.6, 50, 70] and SNR = 500. Panel **(C)**: [0.4, 0.6, 40, 150] and SNR = 20. Panel **(D)**: [0.4, 0.6, 40, 150] and SNR = 500. Panels **(A)** and **(B)**, examples of an ill-posed problem with similar *T*_21_ and *T*_22_, exhibit substantial regions where MSE(*λ*) *<* MSE_Nonreg_. In contrast, panels **(C)** and **(D)** show results for a well-posed case, in which *T*_21_ and *T*_22_ are very different. For **(D)**, there are no values of *λ* for which regularization is helpful. Thus, for greater ill-posedness, **(A)** and **(B)**, there is a greater range of *λ* for which MSE(*λ*) *<* MSE_Nonreg_ than for more well-posed problems, and **(D)**. In panel **(D)**, bCRLB *> MSE*_Nonreg_ in the fourth decimal place; this is an artifact of non-convergence to within this small numerical value of the bCRLB computations after two weeks of machine time on a conventional desktop computer. Finally, in all cases, the values of bCRLB_Nonreg_ and bRCLB_Reg_ for *c*_1_ were less than 2% of the standard CRLB.

**FIGURE 4.**
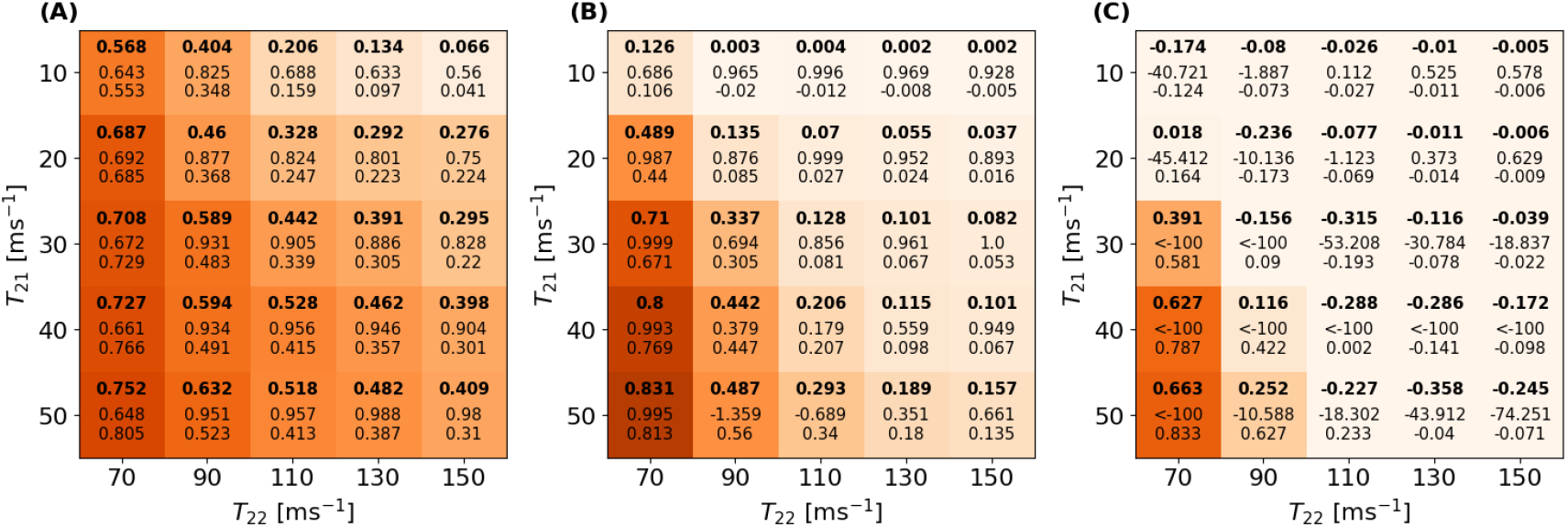
Improvement in MWF Estimation with Regularization. Panel **(A)**: *c*_1_ = 0.2 *c*_2_ = 0.8. Panel **(B)**: *c*_1_ = 0.4 *c*_2_ = 0.6. Panel **(C)**: *c*_1_ = 0.6 *c*_2_ = 0.4. SNR = 20 throughout. Each heat map demonstrates the effect of regularization on the MSE of MWF for the indicated values of *c*_1_ and *c*_2_ across a wide range of *T*_21_ and *T*_22_, more similar values of which, indicating increasing ill-posedness, are found towards the lower-left corner. The three values in each cell are calculated as described in the text and show metrics for the changes in MSE, square bias, and variance with regularization across 500 noise realizations. Results are color-coded for qualitative interpretation, with greater improvements in MSE indicated by deeper orange coloring. As expected, improvements are greater in the more ill-posed regime of parameter space. Further, we see that regularization is of greater value for smaller values of *c*_1_, the surrogate for MWF, reflecting the difficulty in estimation of smaller values of the more rapidly decaying component.

**FIGURE 5.**
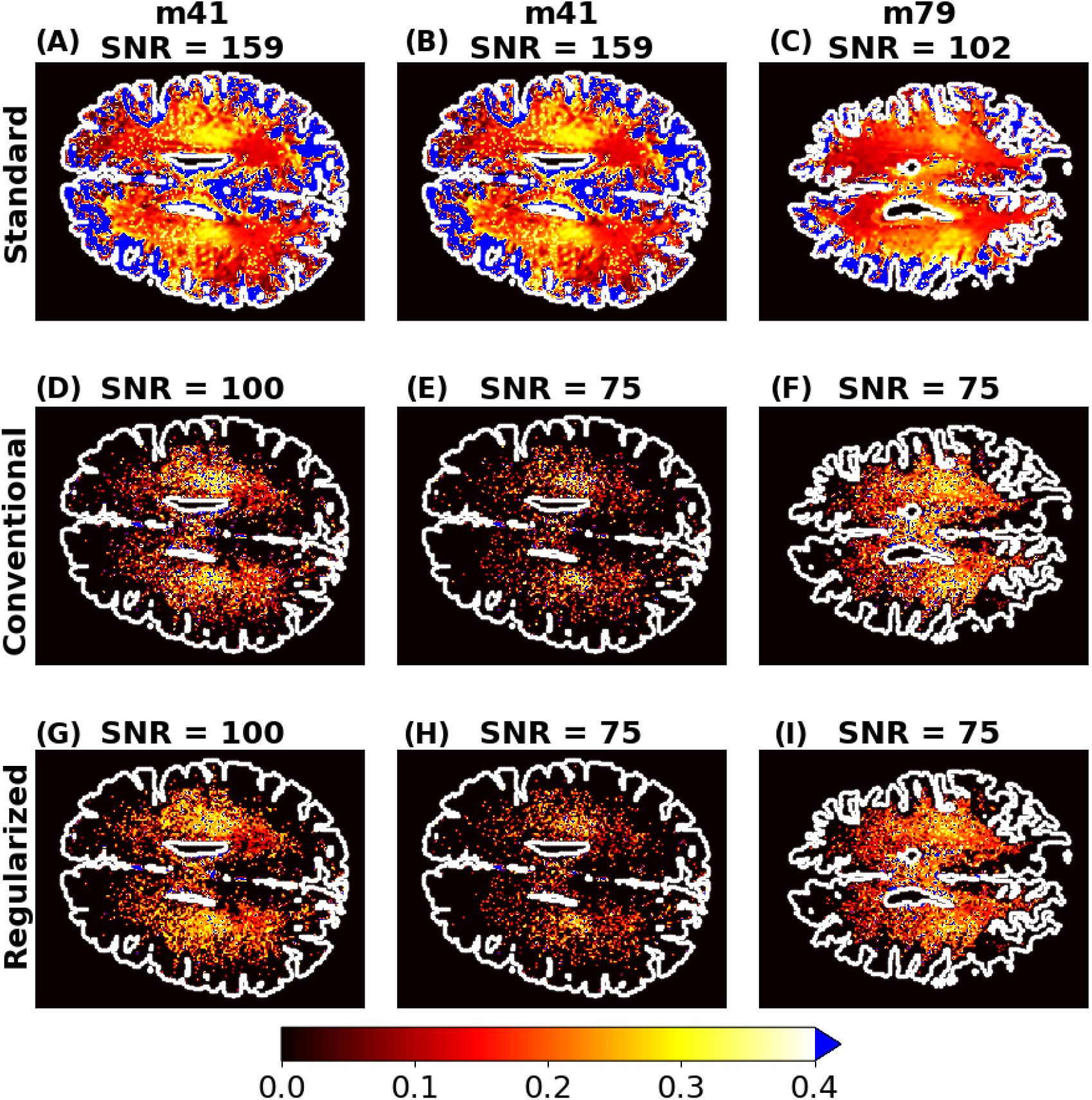
MWF Maps Without and With Regularization. Panels **(A-C)**: Nonregularized MWF map of reference denoised data with SNR = 159 for m41 (repeated in the top row for reference) and SNR = 102 for m79. Panels **(D-F)**: Nonregularized MWF estimates for data with noise added to achieve target SNR values of 100 and 75 for m41, and 75 for m79. Panels **(G-I)**: Regularized MWF estimates corresponding to the nonregularized results in Panels **(D-F)**. Panels **(A, B)**: Mean value of MWF across non-blue pixels = 0.19. Panel **(C)**: Mean value of MWF across pixels in the range of MWF values: 0.18. The color bar shows MWF estimates for pixels found to be biexponential according to the BIC filter. Expected values for MWF are in the range of 0 - 0.4. All voxels greater than 0.4 are colored blue. White is both the outline of the brain and the maximum color bar value. Both visually and in terms of mean MWF value, results from the regularized NLLS analysis are more similar to the reference data than those of the conventional nonregularized analysis.

**FIGURE 6.**
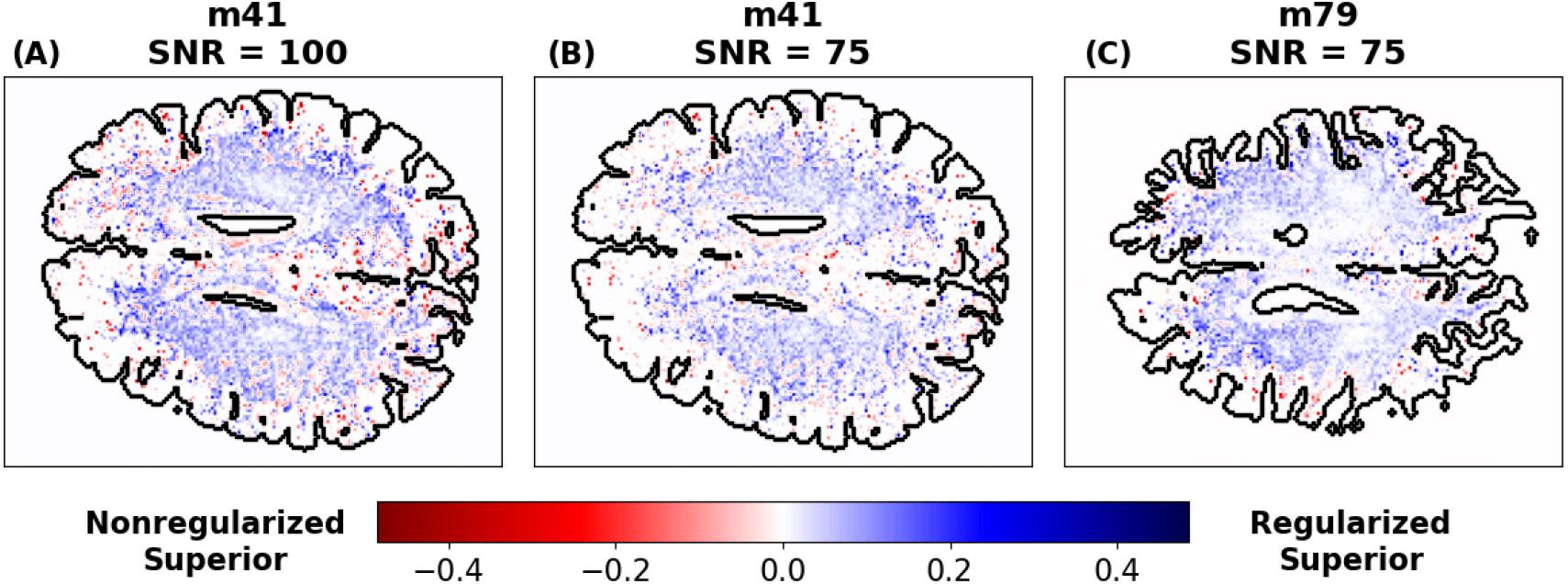
Voxel-wise Improvement in MWF Estimation, ΔRMSE_all,*i*_, Over 20 Noise Realizations. Panel **(A)**: m41, SNR = 100; average across voxels of ΔRMSE_all,*i*_ = 0.025. Panel **(B)**: m41, SNR = 75; average across voxels of ΔRMSE_all,*i*_ = 0.024. Panel **(C)**: m79, SNR = 75; average across voxels of ΔRMSE_all,*i*_ = 0.026. The positive values and dominant blue coloration of the regularized pixels indicate the mean improvement in parameter estimation over multiple noise realizations with regularization.

We evaluated parameter ranges applicable to MWF mapping. Different choices would apply to different tissues [55, 56]. Further, other pulse sequences, including mcDESPOT [57], have also been used for macromolecular mapping; the effect of applying *λ*-NL-RR to these models remains a topic of investigation.

Our simulation analysis was performed for AWGN, the most common noise model in general applications [39]. This is also appropriate for many non-localized MR experiments, and is often used as an approximate model for noise in MRI. Strictly, the noise in magnitude MR imaging is generally Rician or of a more complex type [55, 58]. The Gaussian approximation remains reasonable in the regime of modest SNR [21, 55, 59]. The extension of our work to additional noise models is in progress [55, 59, 60].

Many methods have been developed to select regularization parameters [20, 24, 30]. We found that GCV effectively selected a *λ* that reduced MSE (Figures 2 and 3). Oracle results indicate that further reduction may be possible through improved methods for *λ* selection. Similarly, the performance of *λ*-NL-RR with different regularization penalties remains to be explored [61, 62, 63].

We also applied *λ*-NL-RR to in vivo human brain data. In lieu of gold-standard values, we used parameters estimated from NESMA-filtered images. While we believe these estimates to be accurate [49], the interpretation of our results does not rely strongly upon this assumption. Figure S14 further supports this, showing decreased variance of MWF estimates across noise realizations.

## 5 CONCLUSIONS

We have empirically demonstrated and applied a nonlinear version of the ridge regression theorem to develop a new method for biexponential analysis, *λ*-NL-RR. We found a reduction in MSE through the use of TikReg with *λ* identified by GCV. We applied our analysis to in vivo human brain MRI data for MWF mapping. Our results may have wide applicability to NLLS estimation problems for discrete, unrelated parameters. To our knowledge, regularization has not previously been applied to these ubiquitous parameter estimation problems.

## Abbreviations

MRR: magnetic-resonance relaxometry
LLS: linear least-squares
NLLS: nonlinear least-squares
TikReg: Tikhonov regularization
CRLB: Cramér-Rao Lower Bound
bCRLB: biasedm Cramér-Rao Lower Bound
MWF: myelin water fraction
GCV: generalized cross validation
MSE: mean square error
WM: white matter
GM: grey matter
RRT: ridge regression theorem
RSS: residual sum of squares
BIC: Bayesian information criterion
AWGN: additive white Gaussian noise
CN: condition number

## ACKNOWLEDGMENTS

This work was supported in part by the Intramural Research Program of the National Institute on Aging of the National Institutes of Health.

## SUPPORTING INFORMATION

**Figure S1: Condition Number Comparison of Biexponential and Monoexponential Models**. To illustrate the stability of the biexponential model compared to the monoexponential model, we calculated the condition number (CN) of the Jacobian, *J*, for the biexponential and monoexponential models. *J* was calculated as described in Section 2.1.4 and then evaluated by the “numpy.linalg.cond()” function to determine the CN. For each parameter combination, *J* is unique. Here, we show how variation of one of the transverse decay constants changes the CN. With *T*_22_ free to vary, the other biexponential parameters were fixed at 0.3, 0.7, and 60. The coefficient *c* of the monoexponential model was set at 0.7. As seen, the CN for the monoexponential is less than that for the biexponential. In addition, when *T*_22_ approaches *T*_21_ for the biexponential, the CN rapidly increases indicating the extreme ill-posedness of parameter estimation in this regime.

**Figure S2: Regularized NLLS Objective Function Surface for a Well-Posed Problem**. The regularization parameters are 0, 0.2, and 0.4 for Panels **(A), (B)**, and **(C)** respectively. We generated these regularization surfaces by adding noise to the signal generated by [*c*_1_, *c*_2_, *T*_21_, *T*_22_] = [0.4, 0.6, 40, 150]. Noise was added to achieve an SNR of 100. Subsequent analysis and interpretation is the same as that for Figure 1. For this well-posed case, variance remains small but the effect of regularization to increase bias is clearly seen.

**Figure S3: Improvement in MWF Estimation with Regularization for SNR = 20**. Simulation heat map results for SNR = 20. Panel **(A)**: *c*_1_ = 0.0 *c*_2_ = 1.0. Panel **(B)**: *c*_1_ = 0.05 *c*_2_ = 0.95. Panel **(C)**: *c*_1_ = 0.2 *c*_2_ = 0.8. Panel **(D)**: *c*_1_ = 0.4 *c*_2_ = 0.6. Panel **(E)**: *c*_1_ = 0.5 *c*_2_ = 0.5. Panel **(F)**: *c*_1_ = 0.6 *c*_2_ = 0.4. Panel **(G)**: *c*_1_ = 0.8 *c*_2_ = 0.2. Panel **(H)**: *c*_1_ = 0.95 *c*_2_ = 0.05. Panel **(I)**: *c*_1_ = 1.0 *c*_2_ = 0.0. These results expand upon those in Figure 4.

**Figure S4: Improvement in MWF Estimation with Regularization for SNR = 50**. Simulation heat map results for SNR = 50, with panels otherwise as in Figure S3. Panel **(A)**: *c*_1_ = 0.0 *c*_2_ = 1.0. Panel **(B)**: *c*_1_ = 0.05 *c*_2_ = 0.95. Panel **(C)**: *c*_1_ = 0.2 *c*_2_ = 0.8. Panel **(D)**: *c*_1_ = 0.4 *c*_2_ = 0.6. Panel **(E)**: *c*_1_ = 0.5 *c*_2_ = 0.5. Panel **(F)**: *c*_1_ = 0.6 *c*_2_ = 0.4. Panel **(G)**: *c*_1_ = 0.8 *c*_2_ = 0.2. Panel **(H)**: *c*_1_ = 0.95 *c*_2_ = 0.05. Panel **(I)**: *c*_1_ = 1.0 *c*_2_ = 0.0. Comparing with Figure S3, higher SNR decreases the advantage of regularization.

**Figure S5: Improvement in MWF Estimation with Regularization for SNR = 100**. Simulation heat map results for SNR = 100, with panels otherwise as in Figure S3. Panel **(A)**: *c*_1_ = 0.0 *c*_2_ = 1.0. Panel **(B)**: *c*_1_ = 0.05 *c*_2_ = 0.95. Panel **(C)**: *c*_1_ = 0.2 *c*_2_ = 0.8. Panel **(D)**: *c*_1_ = 0.4 *c*_2_ = 0.6. Panel **(E)**: *c*_1_ = 0.5 *c*_2_ = 0.5. Panel **(F)**: *c*_1_ = 0.6 *c*_2_ = 0.4. Panel **(G)**: *c*_1_ = 0.8 *c*_2_ = 0.2. Panel **(H)**: *c*_1_ = 0.95 *c*_2_ = 0.05. Panel **(I)**: *c*_1_ =1.0 *c*_2_ = 0.0. Higher SNR further decreases the utility of regularization.

**Figure S6: Improvement in MWF Estimation with Regularization for SNR = 500**. Simulation heat map results for SNR = 500, with panels otherwise as in Figure S3. Panel **(A)**: *c*_1_ = 0.0 *c*_2_ = 1.0. Panel **(B)**: *c*_1_ = 0.05 *c*_2_ = 0.95. Panel **(C)**: *c*_1_ = 0.2 *c*_2_ = 0.8. Panel **(D)**: *c*_1_ = 0.4 *c*_2_ = 0.6. Panel **(E)**: *c*_1_ = 0.5 *c*_2_ = 0.5. Panel **(F)**: *c*_1_ = 0.6 *c*_2_ = 0.4. Panel **(G)**: *c*_1_ = 0.8 *c*_2_ = 0.2. Panel **(H)**: *c*_1_ = 0.95 *c*_2_ = 0.05. Panel **(I)**: *c*_1_ = 1.0 *c*_2_ = 0.0. Note that regularization shows little improvement across most (*c*_1_, *c*_2_) combinations at this high SNR.

**Figure S7: BIC Filter Results for Multiple Noise Realizations** As described in the text, the BIC filter approach assigns each voxel signal to either monoexponential or biexponential decay. Panels **(A), (B)**, and **(C)** are for the subjects and SNR as indicated and show the frequency, out of 20 noise realizations, that a pixel is identified as biexponential. As expected, with decreased SNR, the BIC assigns fewer decay curves as biexponential. In all panels, biexponential behavior is most prevalent within the myelinated WM, with the peripheral GM being more frequently identified with monoexponentiality.

**Figure S8: MWF Difference Maps for Single Noise Realizations**. Difference maps corresponding to Figure 5. Panels **(A-C)**: absolute difference between the standard reference NESMA filtered MWF estimates and the nonregularized MWF estimates. Panels **(D-F)**: absolute difference between the standard reference NESMA filtered MWF estimates and the regularized MWF estimates. Panels **(A**,**D)**: m41 with SNR = 100. Panels **(B**,**E)**: m41 with SNR = 75. Panels **(C**,**F)**: m79 with SNR = 75. The white value in the bottom left corner is the average of the absolute difference across voxels where regularization was applied. The improvement from regularization is evident.

**Figure S9: MWF Maps Without and With Regularization for m41**. This figure is analogous to Figure 5 but is applied to the axial slices of m41 for a target SNR of 100. Panels **(A-C)**: Nonregularized MWF map of reference denoised data with SNR = 159 for m41. Panels **(D-F)**: Nonregularized MWF estimates for data at the target SNR value of 100. Panels **(G-I)**: Regularized MWF estimates from the same data used to generate **(D-F)**. Panels **(A**,**D**,**G)**: inferior brain axial slice. Panels **(B**,**E**,**H)**: central brain axial slice. Panels **(C**,**F**,**I)**: superior brain axial slice. Qualitatively, the regularized panels show less fine-scale variability compared to the conventionally estimated MWF results in Panels **(D-F)**. The MWF estimates are in the appropriate range of 0-0.4.

**Figure S10: MWF Difference Maps for Single Noise Realizations of m41**. This figure is analogous to Figure S8 but is applied to the axial slices of m41 for a target SNR of 100. All differences and calculations are done on the same noise realizations from Figure S9 and are specific to different axial slices of m41 with the target SNR of 100. Panels **(A-C)**: absolute difference between the standard reference NESMA filtered MWF estimates and the nonregularized MWF estimates. Panels **(D-F)**: absolute difference between the standard reference NESMA filtered MWF estimates and the regularized MWF estimates. Panels **(A, D)**: an inferior axial slice. Panels **(B, E)**: a central axial slice. Panels **(C, F)**: a superior axial slice. The white value in the bottom left corner is the average value of the absolute differences, only considering voxels where regularization was applied. As in Figure S8, the average absolute difference is less for the regularized MWF estimates than the nonregularized MWF estimates.

**Figure S11: MWF Maps Without and With Regularization for m79**. This figure is analogous to Figure 5 but is applied to the axial slices of m79 for a target SNR of 75. Panels **(A-C)**: Nonregularized MWF map of reference denoised data with SNR = 102 for m79. Panels **(D-F)**: Nonregularized MWF estimates for data at the target SNR value of 75. Panels **(G-I)**: Regularized MWF estimates from the same data to generate **(D-F)**. Panels **(A, D, G)**: inferior brain axial slice. Panels **(B, E, H)**: central brain axial slice. Panels **(C, F, I)**: superior brain axial slice. The same trends and ranges are seen as are observed in Figure S9, with MWF estimates primarily in the range of 0-0.4 and less variability in the regularized result.

**Figure S12: MWF Difference Maps for Single Noise Realizations of m79**. This figure is analogous to Figure S8 but is applied to the axial slices of m79 for a target SNR of 75. All differences and calculations are done on the same noise realizations from Figure S11 and are specific to different axial slices of m79 with the target SNR of 75. Panels **(A-C)**: absolute difference between the standard reference NESMA filtered MWF estimates and the nonregularized MWF estimates. Panels **(D-F)**: absolute difference between the standard reference NESMA filtered MWF estimates and the regularized MWF estimates. Panels **(A, D)**: an inferior axial slice. Panels **(B, E)**: a central axial slice. Panels **(C, F)**: a superior axial slice. The white value in the bottom left corner is the average value of the absolute differences, only considering voxels where regularization was applied. As in Figure S8 and S10, the average absolute difference is less for the regularized MWF estimates than the nonregularized MWF estimates.

**Figure S13: Improvement in MWF Estimation**, Δ**RMSE**_**indiv**_, **for Three Individual Noise Realizations of m41 with SNR = 100**. Panel **(A)**: ΔRMSE_indiv_ = 0.034, an 18% improvement in MSE. Panel **(B)**: ΔRMSE_indiv_ = 0.036, a 19% improvement in MSE. Panel **(C)**: ΔRMSE_indiv_ = 0.033, a 17% improvement in MSE. The positive values and the dominant blue coloration of the regularized pixels in each panel indicate the improvement in parameter estimation with regularization for a single noise realization.

**Figure S14: Effect of Regularization on Standard Deviation in MWF Estimation** Panel **(A)**: ΔSTD_all,*i*_ across 20 noise realizations for a central slice of m41 with a target SNR of 100; the mean value across regularized pixels is 0.034. Panel **(B)**: ΔSTD_all,*i*_ across 20 noise realizations for a central slice of m41 with a target SNR of 75; the mean value across regularized pixels is 0.022. Panel **(C)**: ΔSTD_all,*i*_ across 20 noise realizations for a central slice of m79 with a target SNR of 75; the mean value across regularized pixels is 0.020. The consistent blue pixels and positive ΔSTD_all,*i*_ average value indicate decreased variance with regularization through use of *λ*-NL-RR.

**Figure S15: Effect of Regularization on Absolute Bias in MWF Estimation** Panel **(A)**: Δ|BIAS|_all,*i*_ across 20 noise realizations for the illustrated slice of m41 with a target SNR of 100; the mean value across regularized pixels is 0.003. Panel **(B)**: Δ|BIAS|_all,*i*_ across 20 noise realizations for the illustrated slice of m41 with a target SNR of 75; the mean value across regularized pixels is 0.014. Panel **(C)**: Δ|BIAS|_all,*i*_ across 20 noise realizations for the illustrated slice of m79 with a target SNR of 75; the mean value across regularized pixels is 0.020. The mostly positive values for Δ|BIAS|_all,*i*_ indicate that application of *λ*-NL-RR often reduces bias.

**Figure S16: Voxel-wise Improvement in MWF Estimation for m41**. This figure is analogous to Figure 6 but is applied to the axial slices of m41 for a target SNR of 100. Panel **(A)**: an inferior axial slice with an average ΔRMSE_all,i_ of 0.025. Panel **(B)**: a central axial slice with an average ΔRMSE_all,i_ of 0.025. Panel **(C)**: a superior axial slice with an average ΔRMSE_all,i_ of 0.034. Similar to Figure 6, the dominant blue coloration indicates that regularization consistently reduces the RMSE. **Figure S17: Voxel-wise Improvement in MWF Estimation for m79**. This figure is analogous to Figure 6 but is applied to the axial slices of m79 for a target SNR of 75. Panel **(A)**: an inferior axial slice with an average ΔRMSE_all,i_ of 0.028. Panel **(B)**: a central axial slice with an average ΔRMSE_all,i_ of 0.033. Panel **(C)**: a superior axial slice with an average ΔRMSE_all,i_ of 0.020. As in Figures 6 and S16, the dominant blue coloration indicates that regularization consistently reduces the RMSE.

**Figure S18: Effect of Regularization on Variance**. Panel **(A)**: variance of *c*_1_. Panel **(B)**: variance of *c*_2_. Panel **(C)**: variance of *T*_21_. Panel **(D)**: variance of *T*_22_. To supplement the comparison of MSE values with CRLB measures seen in Figure 2, we present the variance associated with these results. This simplifies the comparison with CRLB and bCRLB measures, although MSE remains the metric of choice. We also plot bCRLB as a function of *λ*, bCRLB(*λ*), providing a comparison reference for var(*λ*). The CRLB and bCRLB measures are unchanged from Figure 2. As expected, the experimental variance, var(*λ*), goes to 0 as *λ* becomes large, as is also seen for bCRLB. Consistent with its definition, bCRLB(*λ*) generally lies below var(*λ*); we attribute the small region where this relationship is violated to non-convergence of the bCRLB(*λ*) calculation, as also seen in Panel **(D)** of Figure 3. Additional parameter relationships are as previously discussed. **Table S1: Summary Metrics of MWF Improvement**. The differences in RMSE due to regularization are collected in this table. For every SNR, slice, and participant combination, the values are positive, indicating that regularization consistently reduces the MSE in estimating MWF. The average ΔRMSE_all,i_ is calculated as the average over ΔRMSE_all,i_ values for each regularized voxel. The average ΔRMSE_indiv_ is calculated as the average of ΔRMSE_indiv_ over 20 noise realizations. The low standard deviation of ΔRMSE_indiv_ values indicate stability of the regularized analysis across noise realizations.

**Table S1: Summary Metrics of MWF Improvement**. The differences in RMSE due to regularization are collected in this table. For every SNR, slice, and participant combination, the values are positive, indicating that regularization consistently reduces the MSE in estimating MWF. The average ΔRMSEall,i is calculated as the average over ΔRMSEall,i values for each regularized voxel. The average ΔRMSEindiv is calculated as the average of ΔRMSEindiv over 20 noise realizations

The low standard deviation of ΔRMSEindiv values indicate stability of the regularized analysis across noise realizations.

**How to cite this article:** Hampton GS, et al (2024), More Accurate Myelin Mapping in the Central Nervous System, *TBD, TBD*.

