## Supplemental Figures and Captions for "Myelin Mapping in the Human Brain Using an Empirical Extension of the Ridge Regression Theorem"

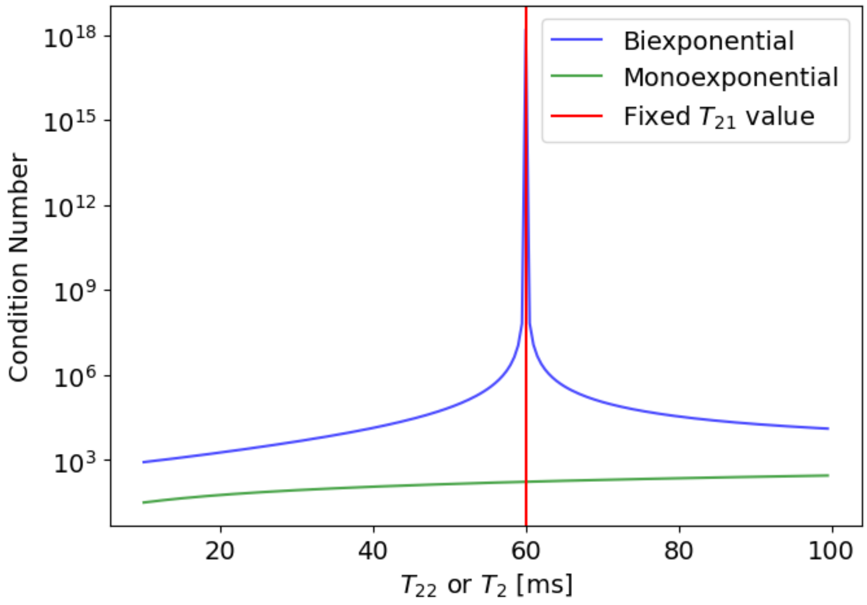


**Figure S1**

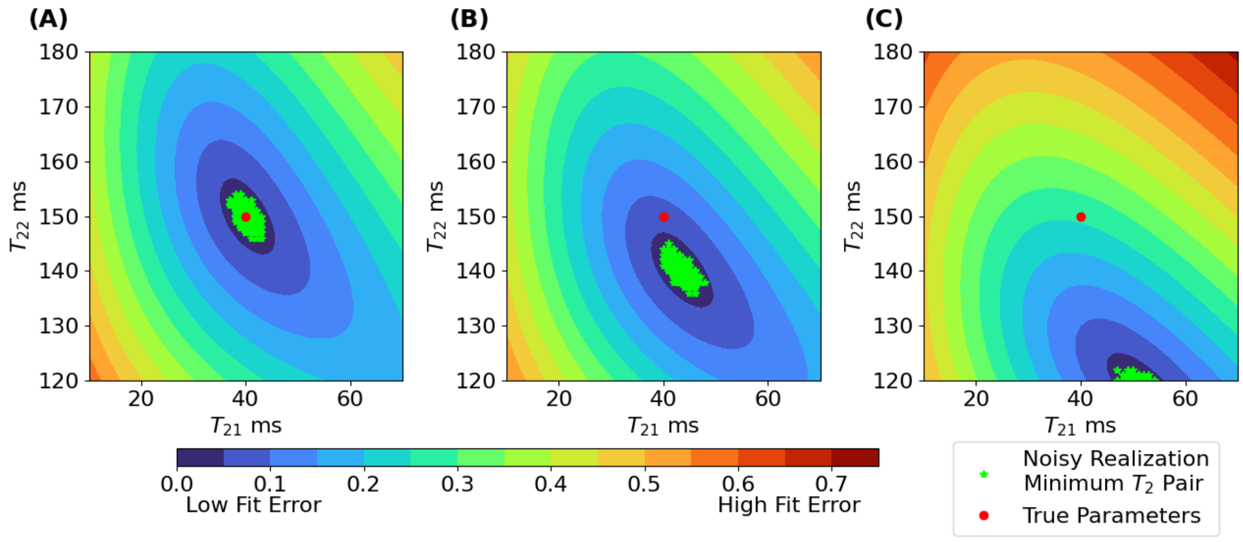


**Figure S2**

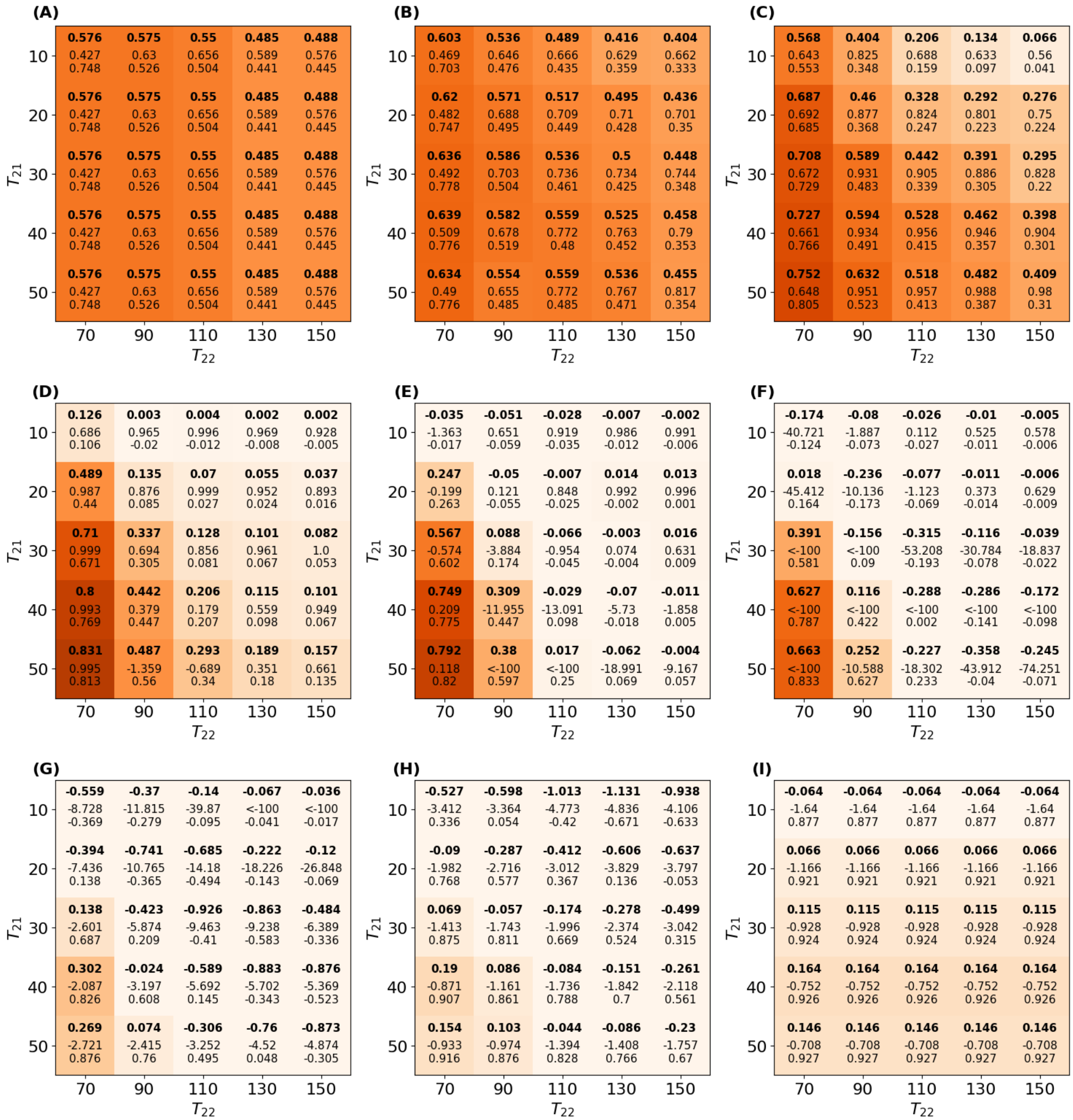
(A)
0.576 0.575 0.55 0.485 0.488
0.427 0.63 0.656 0.589 0.576
0.748 0.526 0.504 0.441 0.445
0.576 0.575 0.55 0.485 0.488
0.427 0.63 0.656 0.589 0.576
0.748 0.526 0.504 0.441 0.445
0.576 0.575 0.55 0.485 0.488
0.427 0.63 0.656 0.589 0.576
0.748 0.526 0.504 0.441 0.445
0.576 0.575 0.55 0.485 0.488
0.427 0.63 0.656 0.589 0.576
0.748 0.526 0.504 0.441 0.445
0.576 0.575 0.55 0.485 0.488
0.427 0.63 0.656 0.589 0.576
0.748 0.526 0.504 0.441 0.445
(B)
0.603 0.536 0.489 0.416 0.404
0.469 0.646 0.666 0.629 0.662
0.703 0.476 0.435 0.359 0.333
0.62 0.571 0.517 0.495 0.436
0.482 0.688 0.709 0.71 0.701
0.747 0.495 0.449 0.428 0.35
0.636 0.586 0.536 0.5 0.448
0.492 0.703 0.736 0.734 0.744
0.778 0.504 0.461 0.425 0.348
0.639 0.582 0.559 0.525 0.458
0.509 0.678 0.772 0.763 0.79
0.776 0.519 0.48 0.452 0.353
0.634 0.554 0.559 0.536 0.455
0.49 0.655 0.772 0.767 0.817
0.776 0.485 0.485 0.471 0.354
(C)
0.568 0.404 0.206 0.134 0.066
0.643 0.825 0.688 0.633 0.56
0.553 0.348 0.159 0.097 0.041
0.687 0.46 0.328 0.292 0.276
0.692 0.877 0.824 0.801 0.75
0.685 0.368 0.247 0.223 0.224
0.708 0.589 0.442 0.391 0.295
0.672 0.931 0.905 0.886 0.828
0.729 0.483 0.339 0.305 0.22
0.727 0.594 0.528 0.462 0.398
0.661 0.934 0.956 0.946 0.904
0.766 0.491 0.415 0.357 0.301
0.752 0.632 0.518 0.482 0.409
0.648 0.951 0.957 0.988 0.98
0.805 0.523 0.413 0.387 0.31
(D)
0.126 0.003 0.004 0.002 0.002
0.686 0.965 0.996 0.969 0.928
0.106 -0.02 -0.012 -0.008 -0.005
0.489 0.135 0.07 0.055 0.037
0.987 0.876 0.999 0.952 0.893
0.44 0.085 0.027 0.024 0.016
0.71 0.337 0.128 0.101 0.082
0.999 0.694 0.856 0.961 1.0
0.671 0.305 0.081 0.067 0.053
0.8 0.442 0.206 0.115 0.101
0.993 0.379 0.179 0.559 0.949
0.769 0.447 0.207 0.098 0.067
0.831 0.487 0.293 0.189 0.157
0.995 -1.359 -0.689 0.351 0.661
0.813 0.56 0.34 0.18 0.135
(E)
-0.035 -0.051 -0.028 -0.007 -0.002
-1.363 0.651 0.919 0.986 0.991
-0.017 -0.059 -0.035 -0.012 -0.006
0.247 -0.05 -0.007 0.014 0.013
-0.199 0.121 0.848 0.992 0.996
0.263 -0.055 -0.025 -0.002 0.001
0.567 0.088 -0.066 -0.003 0.016
-0.574 -3.884 -0.954 0.074 0.631
0.602 0.174 -0.045 -0.004 0.009
0.749 0.309 -0.029 -0.07 -0.011
0.209 -11.955 -13.091 -5.73 -1.858
0.775 0.447 0.098 -0.018 0.005
0.792 0.38 0.017 -0.062 -0.004
0.118 <-100 <-100 -18.991 -9.167
0.82 0.597 0.25 0.069 0.057
(F)
-0.174 -0.08 -0.026 -0.01 -0.005
-40.721 -1.887 0.112 0.525 0.578
-0.124 -0.073 -0.027 -0.011 -0.006
0.018 -0.236 -0.077 -0.011 -0.006
-45.412 -10.136 -1.123 0.373 0.629
0.164 -0.173 -0.069 -0.014 -0.009
0.391 -0.156 -0.315 -0.116 -0.039
<-100 <-100 -53.208 -30.784 -18.837
0.581 0.09 -0.193 -0.078 -0.022
0.627 0.116 -0.288 -0.286 -0.172
<-100 <-100 <-100 <-100 <-100
0.787 0.422 0.002 -0.141 -0.098
0.663 0.252 -0.227 -0.358 -0.245
<-100 -10.588 -18.302 -43.912 -74.251
0.833 0.627 0.233 -0.04 -0.071
(G)
-0.559 -0.37 -0.14 -0.067 -0.036
-8.728 -11.815 -39.87 <-100 <-100
-0.369 -0.279 -0.095 -0.041 -0.017
-0.394 -0.741 -0.685 -0.222 -0.12
-7.436 -10.765 -14.18 -18.226 -26.848
0.138 -0.365 -0.494 -0.143 -0.069
0.138 -0.423 -0.926 -0.863 -0.484
-2.601 -5.874 -9.463 -9.238 -6.389
0.687 0.209 -0.41 -0.583 -0.336
0.302 -0.024 -0.589 -0.883 -0.876
-2.087 -3.197 -5.692 -5.702 -5.369
0.826 0.608 0.145 -0.343 -0.523
0.269 0.074 -0.306 -0.76 -0.873
-2.721 -2.415 -3.252 -4.52 -4.874
0.876 0.76 0.495 0.048 -0.305
(H)
-0.527 -0.598 -1.013 -1.131 -0.938
-3.412 -3.364 -4.773 -4.836 -4.106
0.336 0.054 -0.42 -0.671 -0.633
-0.09 -0.287 -0.412 -0.606 -0.637
-1.982 -2.716 -3.012 -3.829 -3.797
0.768 0.577 0.367 0.136 -0.053
0.069 -0.057 -0.174 -0.278 -0.499
-1.413 -1.743 -1.996 -2.374 -3.042
0.875 0.811 0.669 0.524 0.315
0.19 0.086 -0.084 -0.151 -0.261
-0.871 -1.161 -1.736 -1.842 -2.118
0.907 0.861 0.788 0.7 0.561
0.154 0.103 -0.044 -0.086 -0.23
-0.933 -0.974 -1.394 -1.408 -1.757
0.916 0.876 0.828 0.766 0.67
(I)
-0.064 -0.064 -0.064 -0.064 -0.064
-1.64 -1.64 -1.64 -1.64 -1.64
0.877 0.877 0.877 0.877 0.877
0.066 0.066 0.066 0.066 0.066
-1.166 -1.166 -1.166 -1.166 -1.166
0.921 0.921 0.921 0.921 0.921
0.115 0.115 0.115 0.115 0.115
-0.928 -0.928 -0.928 -0.928 -0.928
0.924 0.924 0.924 0.924 0.924
0.164 0.164 0.164 0.164 0.164
-0.752 -0.752 -0.752 -0.752 -0.752
0.926 0.926 0.926 0.926 0.926
0.146 0.146 0.146 0.146 0.146
-0.708 -0.708 -0.708 -0.708 -0.708
0.927 0.927 0.927 0.927 0.927
$T_{21}$
10 20 30 40 50
$T_{22}$
70 90 110 130 150

**Figure S3**

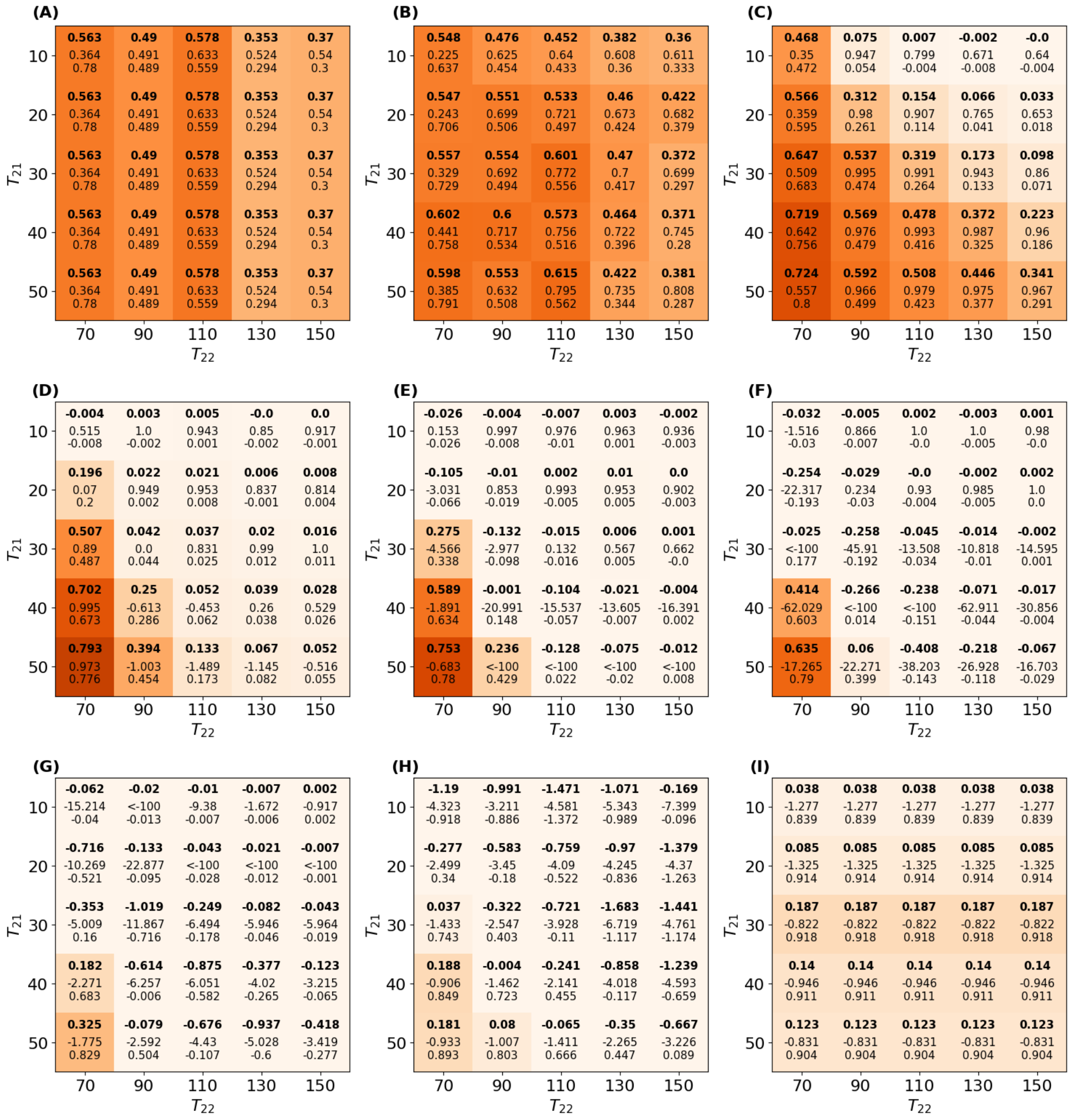

**Figure S4**

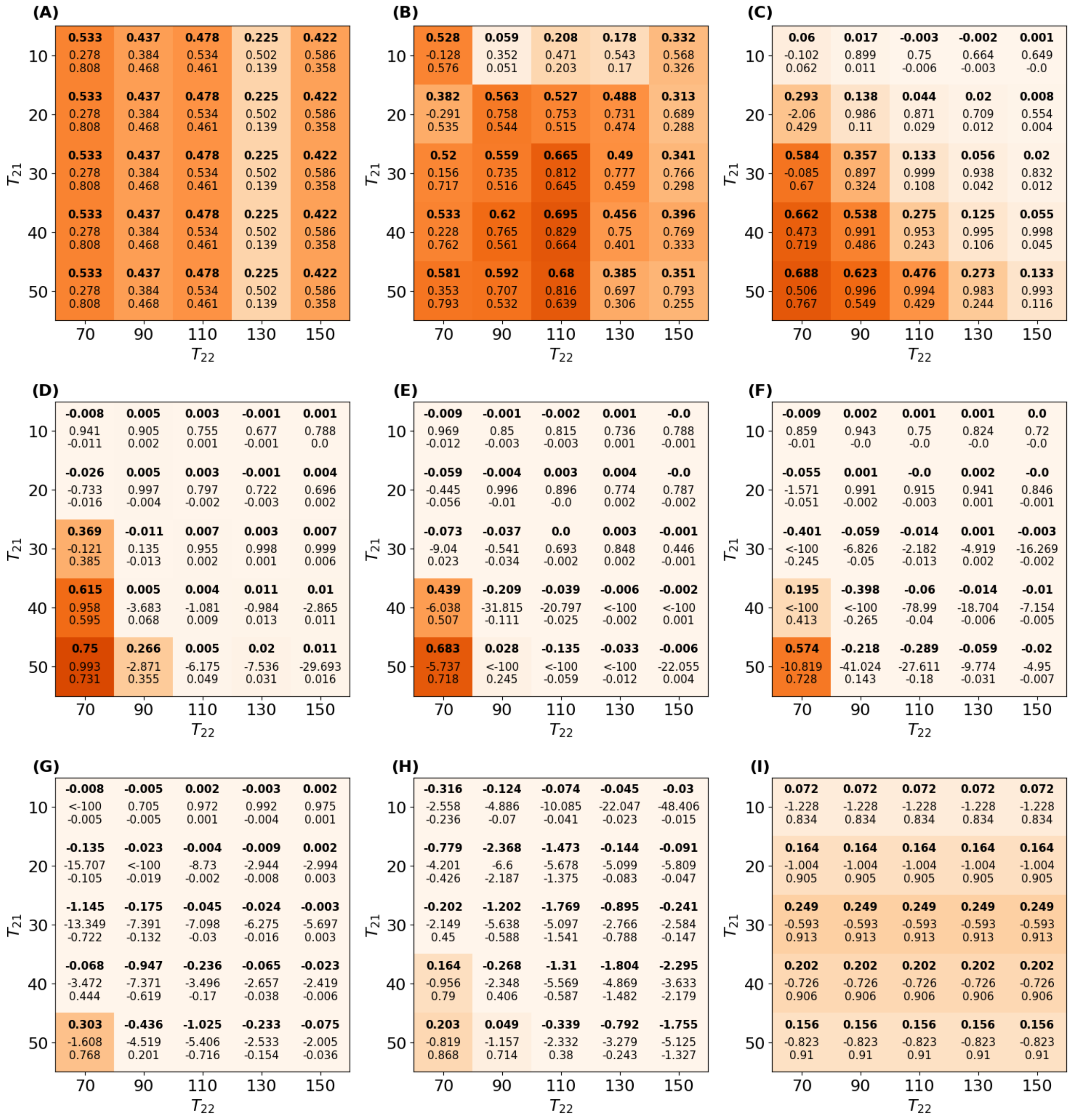

(A)
0.533 0.278 0.808
0.437 0.384 0.468
0.478 0.534 0.461
0.225 0.502 0.139
0.422 0.586 0.358
0.533 0.278 0.808
0.437 0.384 0.468
0.478 0.534 0.461
0.225 0.502 0.139
0.422 0.586 0.358
0.533 0.278 0.808
0.437 0.384 0.468
0.478 0.534 0.461
0.225 0.502 0.139
0.422 0.586 0.358
0.533 0.278 0.808
0.437 0.384 0.468
0.478 0.534 0.461
0.225 0.502 0.139
0.422 0.586 0.358
0.533 0.278 0.808
0.437 0.384 0.468
0.478 0.534 0.461
0.225 0.502 0.139
0.422 0.586 0.358
(B)
0.528 -0.128 0.576
0.059 0.352 0.051
0.208 0.471 0.203
0.178 0.543 0.17
0.332 0.568 0.326
0.382 -0.291 0.535
0.563 0.758 0.544
0.527 0.753 0.515
0.488 0.731 0.474
0.313 0.689 0.288
0.52 0.156 0.717
0.559 0.735 0.516
0.665 0.812 0.645
0.49 0.777 0.459
0.341 0.766 0.298
0.533 0.228 0.762
0.62 0.765 0.561
0.695 0.829 0.664
0.456 0.75 0.401
0.396 0.769 0.333
0.581 0.353 0.793
0.592 0.707 0.532
0.68 0.816 0.639
0.385 0.697 0.306
0.351 0.793 0.255
(C)
0.06 -0.102 0.062
0.017 0.899 0.011
-0.003 0.75 -0.006
-0.002 0.664 -0.003
0.001 0.649 -0.0
0.293 -2.06 0.429
0.138 0.986 0.11
0.044 0.871 0.029
0.02 0.709 0.012
0.008 0.554 0.004
0.584 -0.085 0.67
0.357 0.897 0.324
0.133 0.999 0.108
0.056 0.938 0.042
0.02 0.832 0.012
0.662 0.473 0.719
0.538 0.991 0.486
0.275 0.953 0.243
0.125 0.995 0.106
0.055 0.998 0.045
0.688 0.506 0.767
0.623 0.996 0.549
0.476 0.994 0.429
0.273 0.983 0.244
0.133 0.993 0.116
(D)
-0.008 0.941 -0.011
0.005 0.905 0.002
0.003 0.755 0.001
-0.001 0.677 -0.001
0.001 0.788 0.0
-0.026 -0.733 -0.016
0.005 0.997 -0.004
0.003 0.797 -0.002
-0.001 0.722 -0.003
0.004 0.696 0.002
0.369 -0.121 0.385
-0.011 0.135 -0.013
0.007 0.955 0.002
0.003 0.998 0.001
0.007 0.999 0.006
0.615 0.958 0.595
0.005 -3.683 0.068
0.004 -1.081 0.009
0.011 -0.984 0.013
0.01 -2.865 0.011
0.75 0.993 0.731
0.266 -2.871 0.355
0.005 -6.175 0.049
0.02 -7.536 0.031
0.011 -29.693 0.016
(E)
-0.009 0.969 -0.012
-0.001 0.85 -0.003
-0.002 0.815 -0.003
0.001 0.736 0.001
-0.0 0.788 -0.001
-0.059 -0.445 -0.056
-0.004 0.996 -0.01
0.003 0.896 -0.0
0.004 0.774 0.002
-0.0 0.787 -0.002
-0.073 -9.04 0.023
-0.037 -0.541 -0.034
0.0 0.693 -0.002
0.003 0.848 0.002
-0.001 0.446 -0.001
0.439 -6.038 0.507
-0.209 -31.815 -0.111
-0.039 -20.797 -0.025
-0.006 <-100 -0.002
-0.002 <-100 0.001
0.683 -5.737 0.718
0.028 <-100 0.245
-0.135 <-100 -0.059
-0.033 <-100 -0.012
-0.006 -22.055 0.004
(F)
-0.009 0.859 -0.01
0.002 0.943 -0.0
0.001 0.75 -0.0
0.001 0.824 -0.0
0.0 0.72 -0.0
-0.055 -1.571 -0.051
0.001 0.991 -0.002
-0.0 0.915 -0.003
0.002 0.941 0.001
-0.0 0.846 -0.001
-0.401 <-100 -0.245
-0.059 -6.826 -0.05
-0.014 -2.182 -0.013
0.001 -4.919 0.002
-0.003 -16.269 -0.002
0.195 <-100 0.413
-0.398 <-100 -0.265
-0.06 -78.99 -0.04
-0.014 -18.704 -0.006
-0.01 -7.154 -0.005
0.574 -10.819 0.728
-0.218 -41.024 0.143
-0.289 -27.611 -0.18
-0.059 -9.774 -0.031
-0.02 -4.95 -0.007
(G)
-0.008 <-100 -0.005
-0.005 0.705 -0.005
0.002 0.972 0.001
-0.003 0.992 -0.004
0.002 0.975 0.001
-0.135 -15.707 -0.105
-0.023 <-100 -0.019
-0.004 -8.73 -0.002
-0.009 -2.944 -0.008
0.002 -2.994 0.003
-1.145 -13.349 -0.722
-0.175 -7.391 -0.132
-0.045 -7.098 -0.03
-0.024 -6.275 -0.016
-0.003 -5.697 0.003
-0.068 -3.472 0.444
-0.947 -7.371 -0.619
-0.236 -3.496 -0.17
-0.065 -2.657 -0.038
-0.023 -2.419 -0.006
0.303 -1.608 0.768
-0.436 -4.519 0.201
-1.025 -5.406 -0.716
-0.233 -2.533 -0.154
-0.075 -2.005 -0.036
(H)
-0.316 -2.558 -0.236
-0.124 -4.886 -0.07
-0.074 -10.085 -0.041
-0.045 -22.047 -0.023
-0.03 -48.406 -0.015
-0.779 -4.201 -0.426
-2.368 -6.6 -2.187
-1.473 -5.678 -1.375
-0.144 -5.099 -0.083
-0.091 -5.809 -0.047
-0.202 -2.149 0.45
-1.202 -5.638 -0.588
-1.769 -5.097 -1.541
-0.895 -2.766 -0.788
-0.241 -2.584 -0.147
0.164 -0.956 0.79
-0.268 -2.348 0.406
-1.31 -5.569 -0.587
-1.804 -4.869 -1.482
-2.295 -3.633 -2.179
0.203 -0.819 0.868
0.049 -1.157 0.714
-0.339 -2.332 0.38
-0.792 -3.279 -0.243
-1.755 -5.125 -1.327
(I)
0.072 -1.228 0.834
0.072 -1.228 0.834
0.072 -1.228 0.834
0.072 -1.228 0.834
0.072 -1.228 0.834
0.164 -1.004 0.905
0.164 -1.004 0.905
0.164 -1.004 0.905
0.164 -1.004 0.905
0.164 -1.004 0.905
0.249 -0.593 0.913
0.249 -0.593 0.913
0.249 -0.593 0.913
0.249 -0.593 0.913
0.249 -0.593 0.913
0.202 -0.726 0.906
0.202 -0.726 0.906
0.202 -0.726 0.906
0.202 -0.726 0.906
0.202 -0.726 0.906
0.156 -0.823 0.91
0.156 -0.823 0.91
0.156 -0.823 0.91
0.156 -0.823 0.91
0.156 -0.823 0.91
T21
10
20
30
40
50
T22
70
90
110
130
150


**Figure S5**

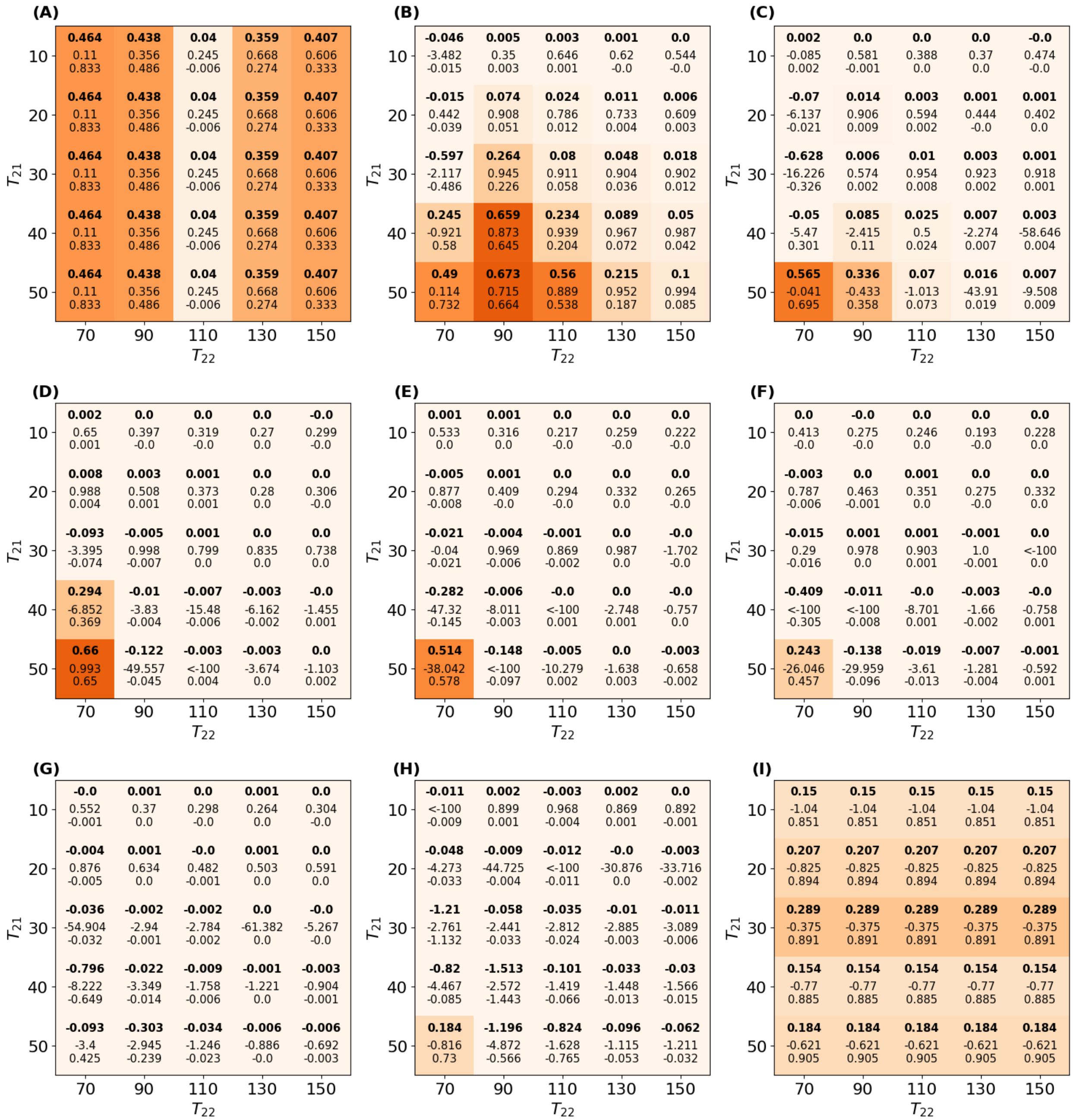


**Figure S6**

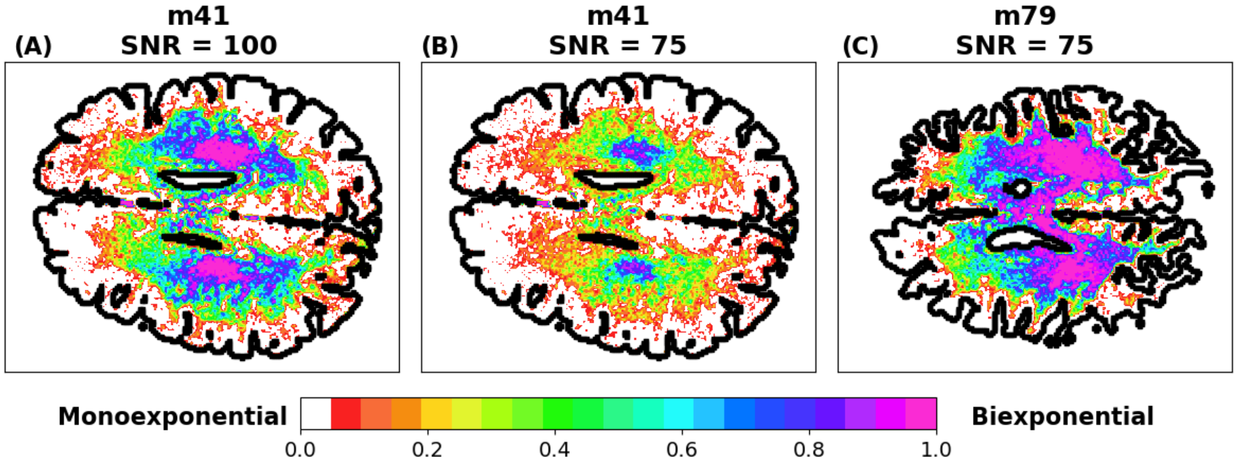


**Figure S7**

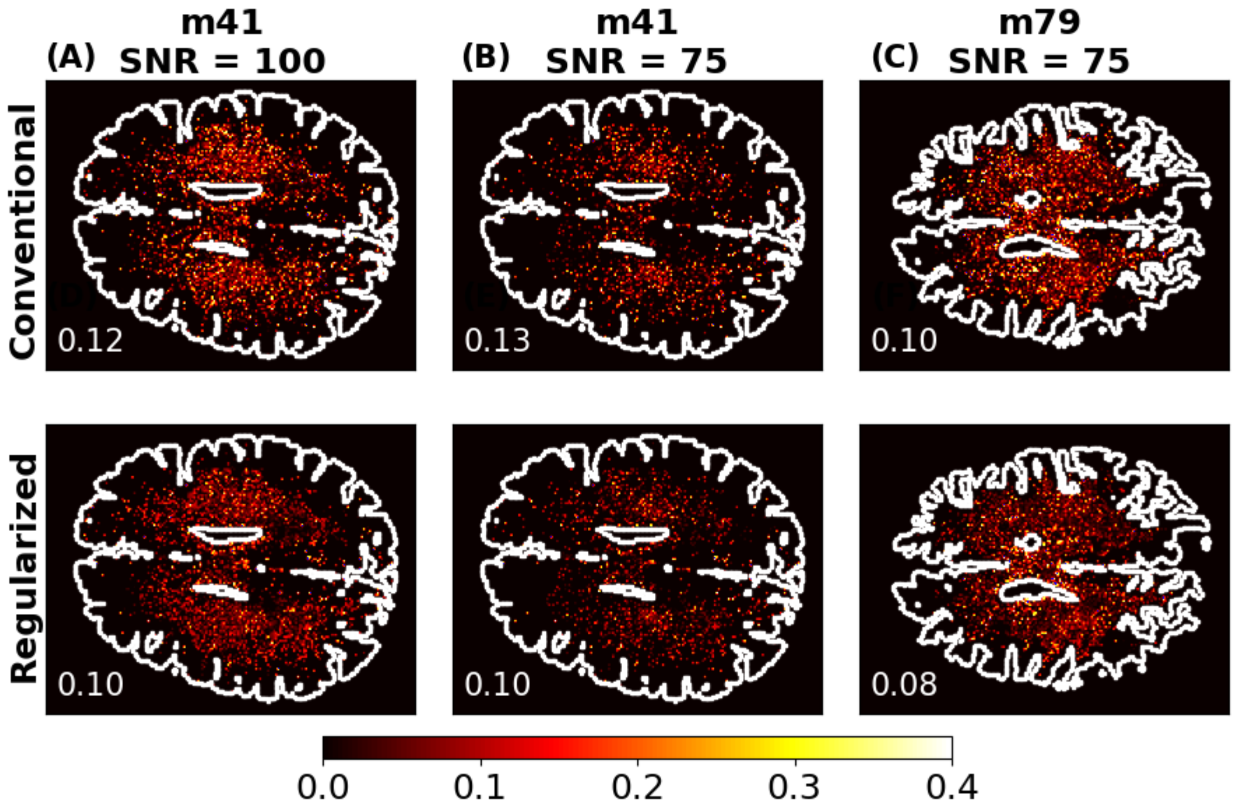


**Figure S8**

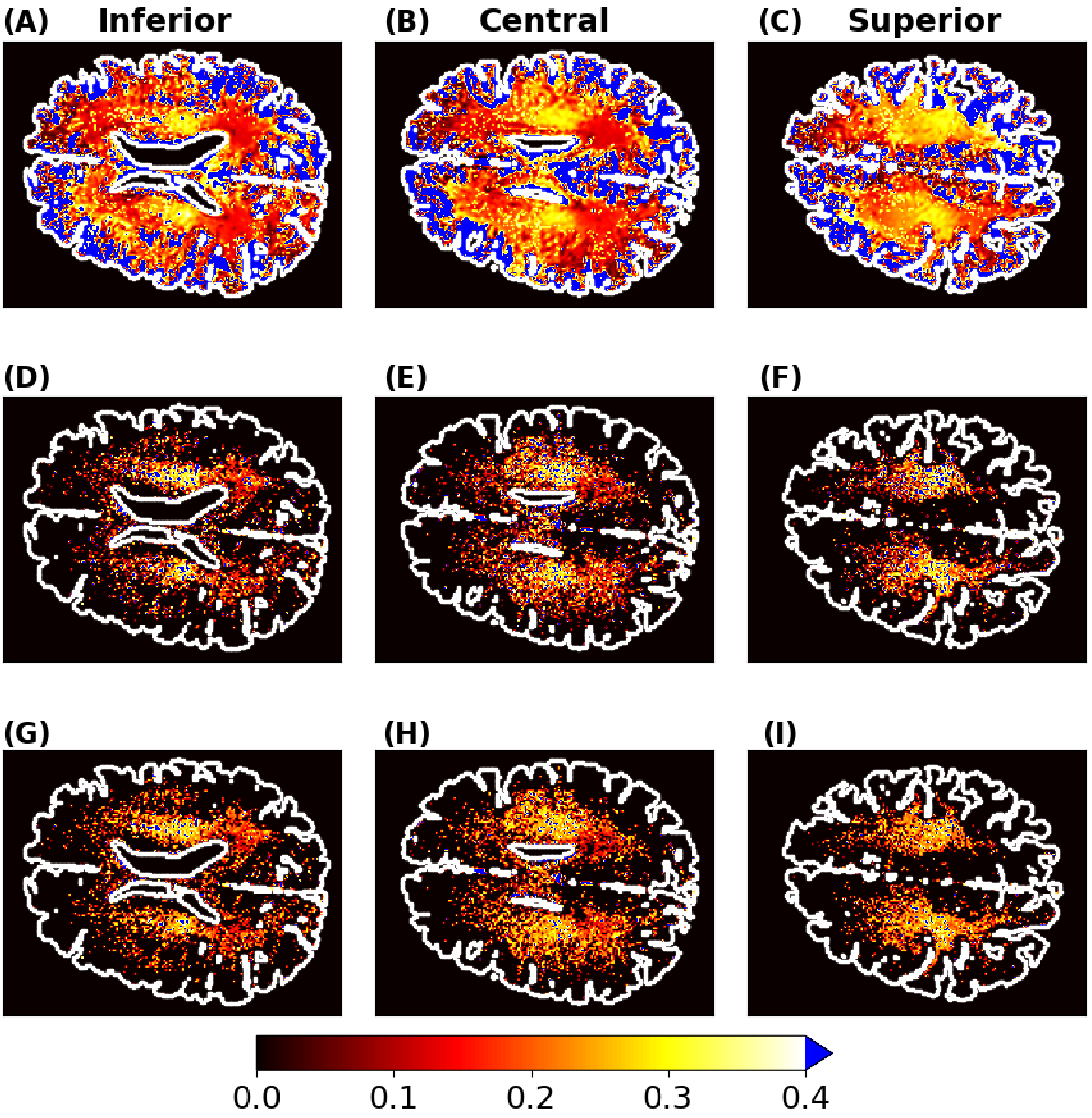


**Figure S9**

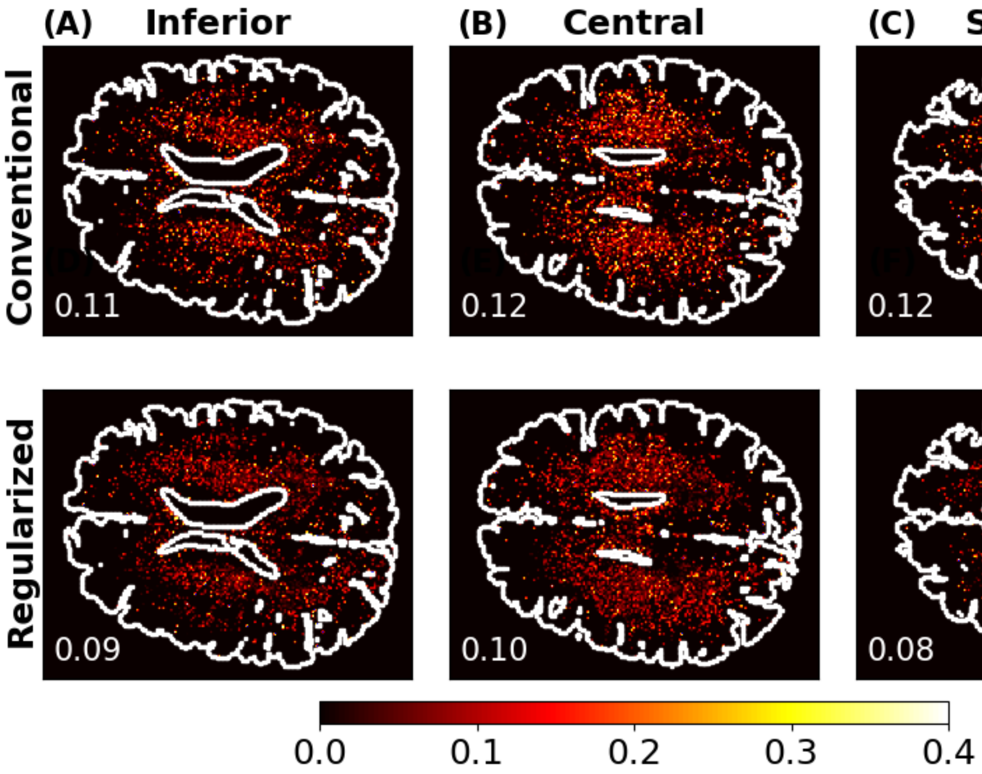

(A) Inferior
(B) Central
(C) Superior
Conventional
Regularized
0.11
0.12
0.12
0.09
0.10
0.08
0.0
0.1
0.2
0.3
0.4


**Figure S10**

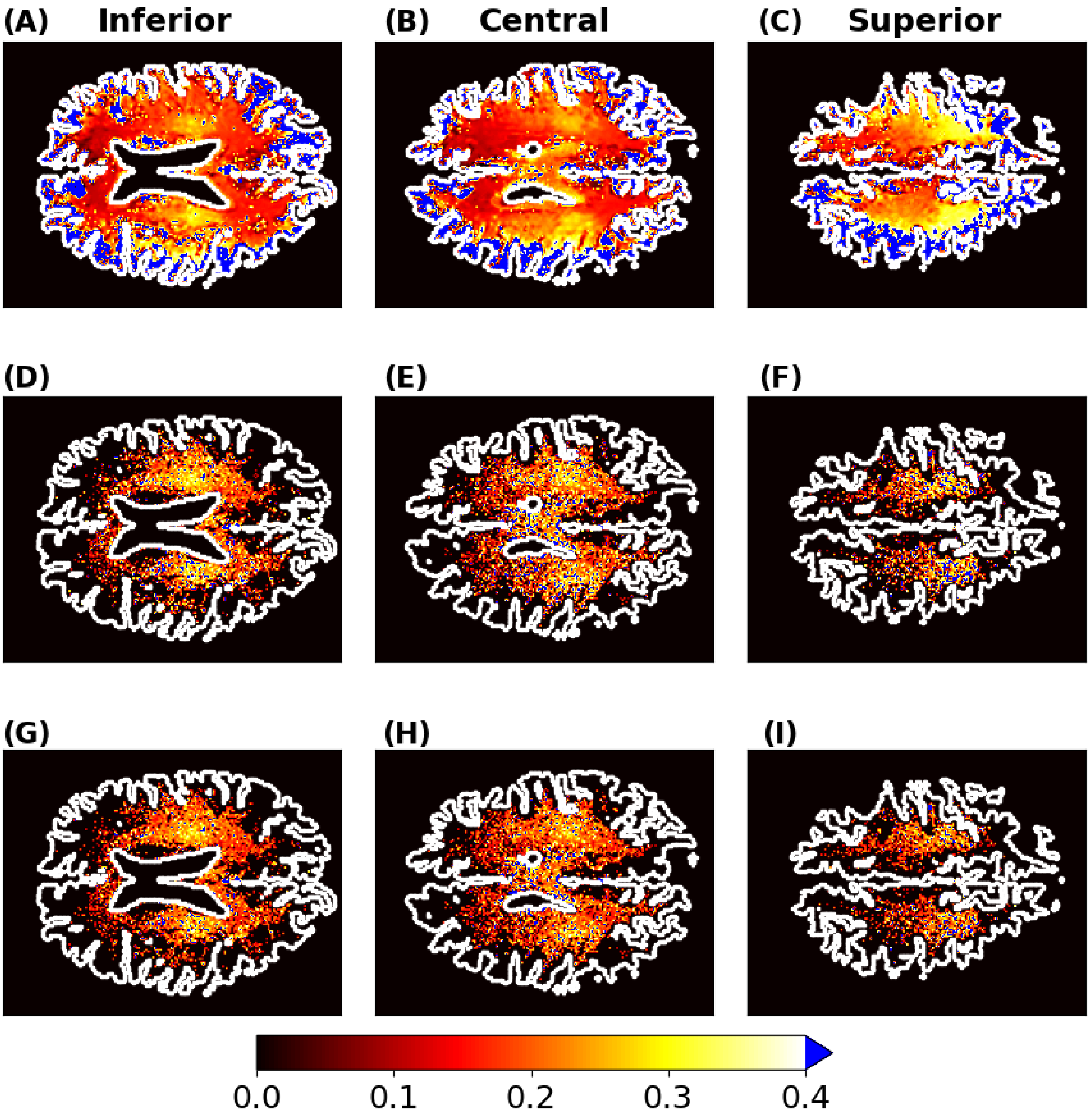


**Figure S11**

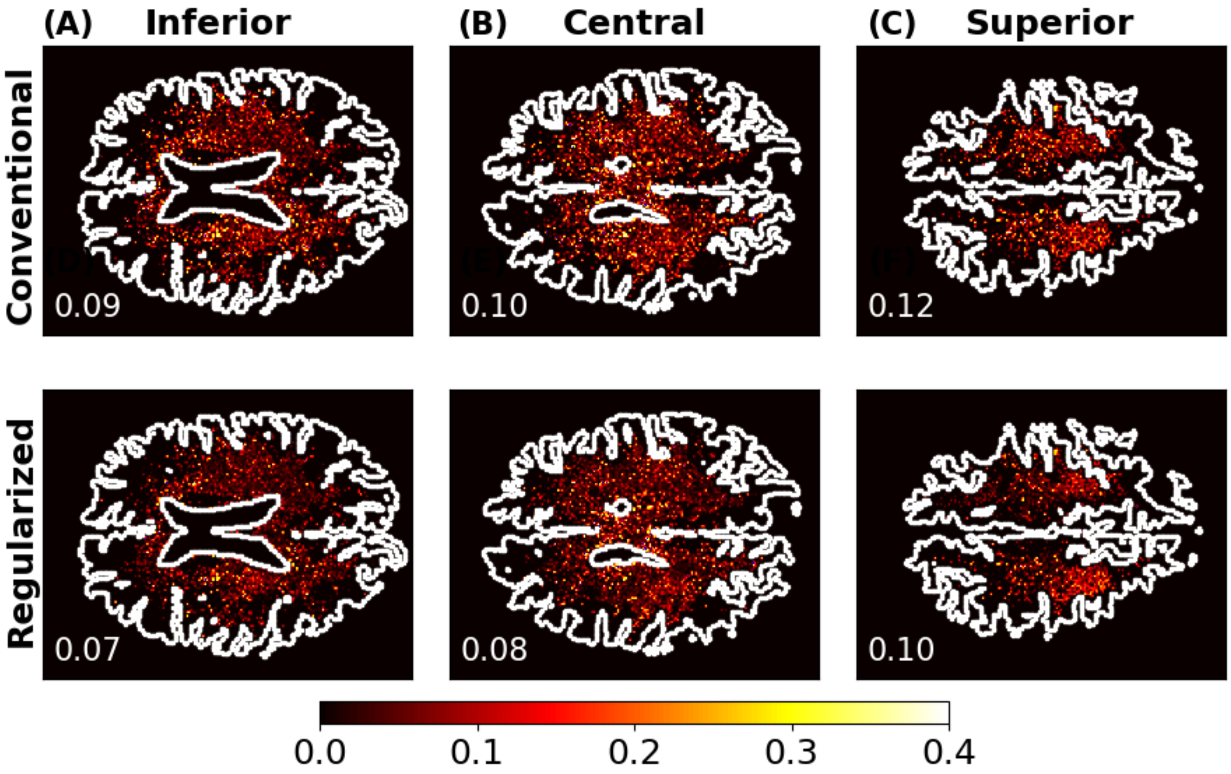
(A) Inferior
(B) Central
(C) Superior
Conventional
Regularized
0.09
0.10
0.12
0.07
0.08
0.10
0.0
0.1
0.2
0.3
0.4

**Figure S12**

The effectiveness for the $i^{\text{th}}$ voxel in a single noise realization was reduced to $\Delta\text{RMSE}_{voxel,i}$ defined in Eq. S1.

$$\Delta\text{RMSE}_{voxel,i} = \left|c_{1,NR,i} - c_{1,i}\right| - \left|c_{1,\text{Reg},i} - c_{1,i}\right|$$

where $c_{1,i}$ is the gold standard value and $c_{1,NR,i}$ and $c_{1,\text{Reg},i}$ are the estimates obtained for non-regularized and regularized NLLS, respectively.

To evaluate all voxels across an individual noise realization, we calculated the $\Delta\text{RMSE}_{\text{indiv}}$ defined in Eq. S2.

$$\Delta\text{RMSE}_{\text{indiv}} = \sqrt{\frac{1}{n}\sum_{i=1}^{n}\left(c_{1,NR,i} - c_{1,i}\right)^2} - \sqrt{\frac{1}{n}\sum_{i=1}^{n}\left(c_{1,\text{Reg},i} - c_{1,i}\right)^2}$$

where n is the number of voxels undergoing regularization, that is, assigned to a biexponential decay by the BIC.

Comparison of conventional, non-regularized to regularized $c_1$ parameter estimates for a single noise realization are shown in Figure S6. WM regions were assigned to monoexponential decay, so that no regularization was applied. Blue and red regions indicate pixels where the decay was characterized as biexponential, with blue showing improvement with regularization and red showing worsening MSE with regularization as defined by Eq. S1.

A summary metric for a noise realization was calculated using Eq. S2, quantifying the change in MSE across all voxels. For the typical noise realization shown, $\Delta\text{RMSE}_{\text{indiv}}$ = 0.034, indicating overall improvement (decrease) in MSE with regularization. Note that the overall average of $c_1$ representing MWF is approximately 0.19, corresponding to an improvement of 13\%. Applying this same analysis to all 20 noise realizations, 20 $\Delta\text{RMSE}_{\text{indiv}}$ values are generated. We calculate the mean, standard deviation, minimum and maximum values for the $\Delta\text{RMSE}_{\text{indiv}}$ values for m41 and m79 at different scans levels in Table S1. We also include the average $\Delta\text{RMSE}_{voxel,i}$ for comparison in Table S1.

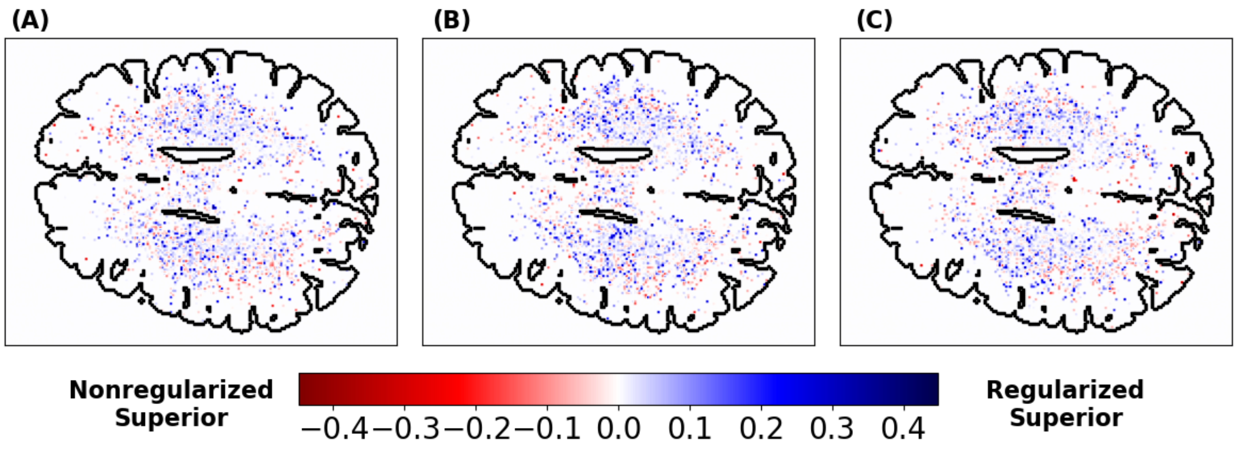


**Figure S13**

We calculated comparable metrics to $\Delta \mathrm{RMSE}_{\mathrm{all},i}$ for the changes in absolute bias, $\Delta|\mathrm{BIAS}|_{\mathrm{all},i}$, and standard deviation, $\Delta \mathrm{STD}_{\mathrm{all},i}$. We define $\overline{c_{1,\mathrm{NR},i}} = \frac{1}{m}\sum_{j\in m} c_{1,\mathrm{NR},i,j}$ and, analogously, $\overline{c_{1,\mathrm{Reg},i}} = \frac{1}{m}\sum_{j\in m} c_{1,\mathrm{Reg},i,j}$ for the $j^{th}$ of $m$ noise realizations where the data for the $i^{th}$ voxel is evaluated as biexponential by the BIC. We have

$$\Delta|\mathrm{BIAS}|_{\mathrm{all},i} = \sqrt{\left(\overline{c_{1,\mathrm{NR},i}} - c_{1,i}\right)^2} - \sqrt{\left(\overline{c_{1,\mathrm{Reg},i}} - c_{1,i}\right)^2}$$

and

$$\Delta \mathrm{STD}_{\mathrm{all},i} = \sqrt{\frac{1}{m}\sum_{j\in m}\left(c_{1,\mathrm{NR},i,j} - \overline{c_{1,\mathrm{NR},i}}\right)^2} - \sqrt{\frac{1}{m}\sum_{j\in m}\left(c_{1,\mathrm{Reg},i,j} - \overline{c_{1,\mathrm{Reg},i}}\right)^2}.$$

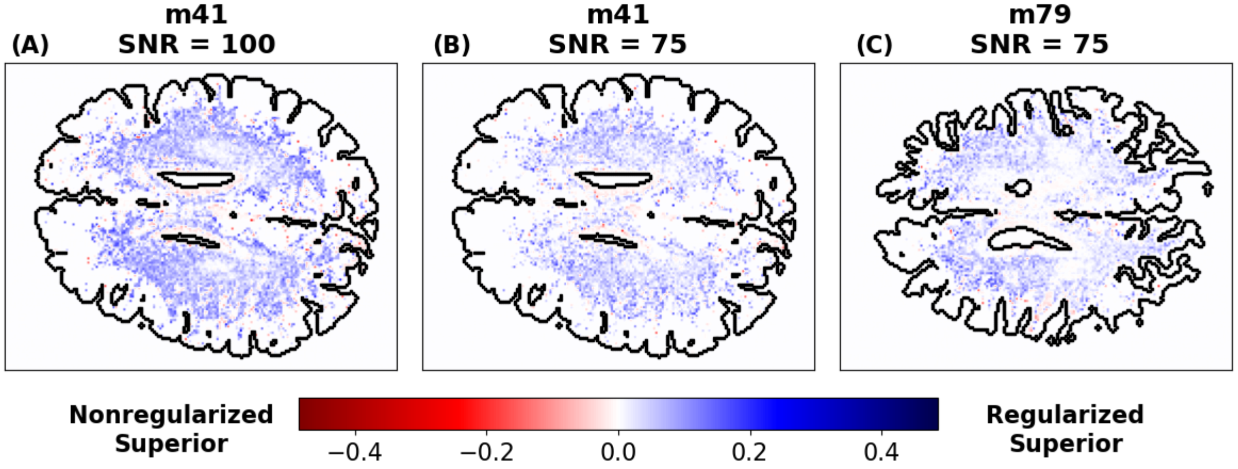


**Figure S14**

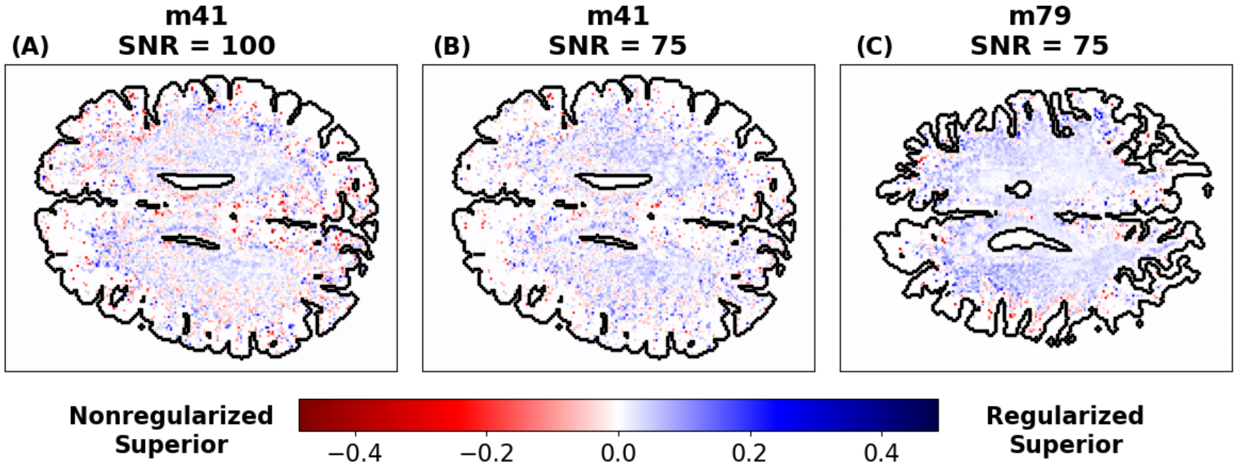


**Figure S15**

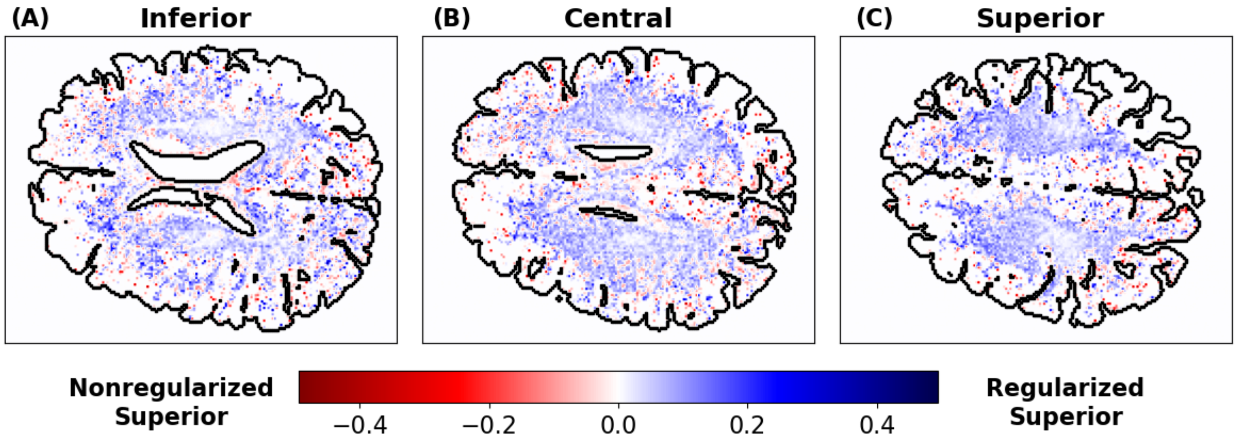


**Figure S16**

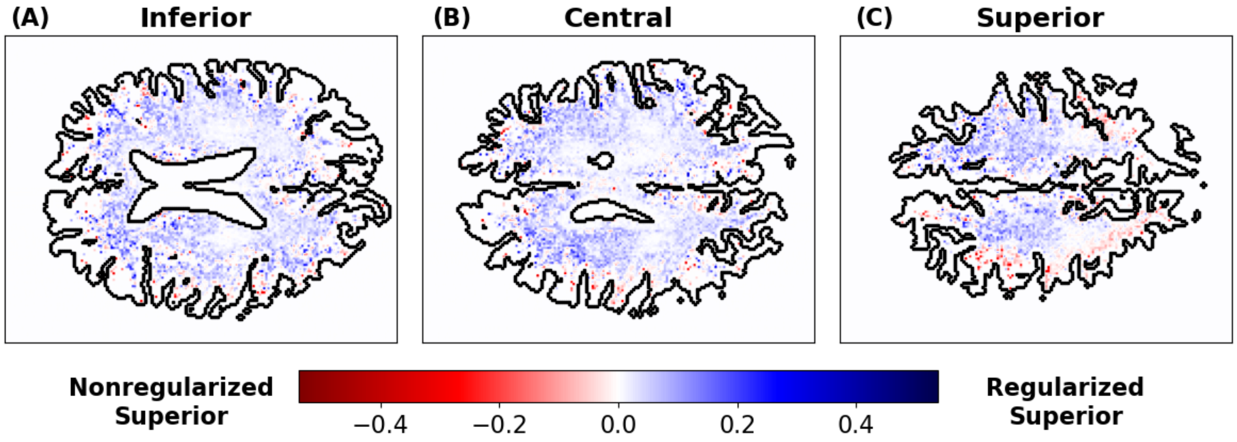


**Figure S17**

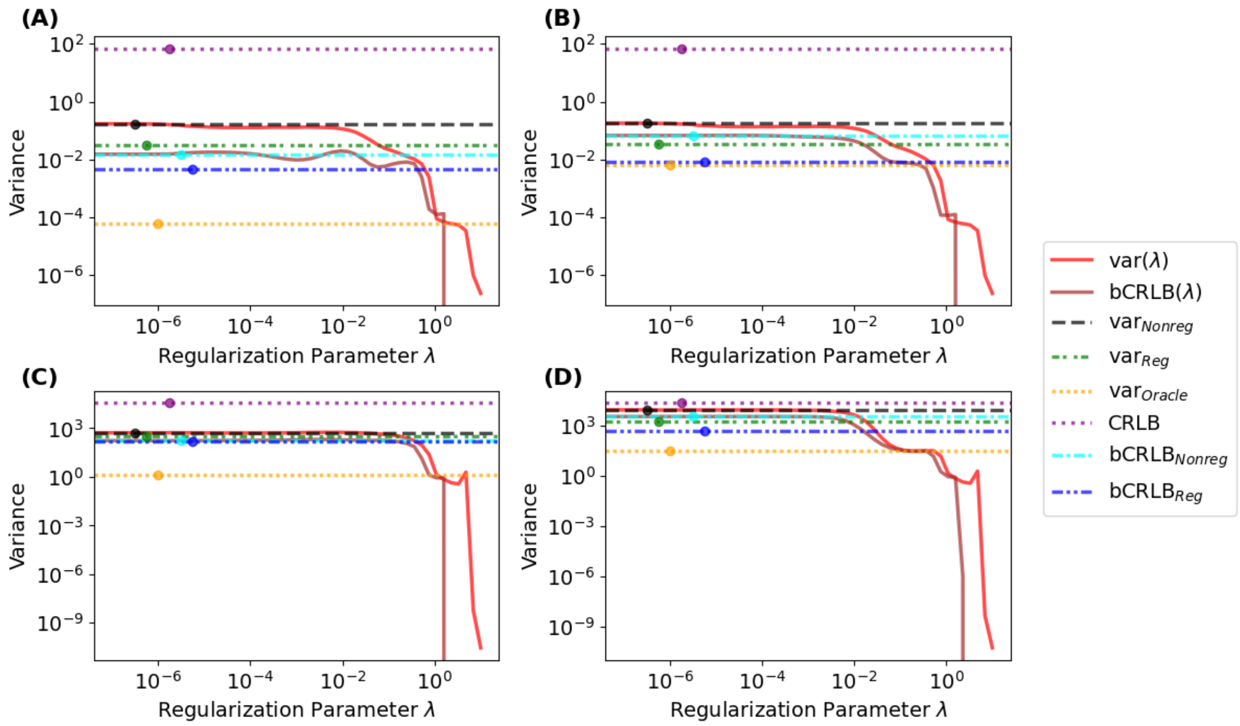


**Figure S18**

| Participant | Target SNR | Scan Level | Average $\Delta\mathrm{RMSE}_{\mathrm{all},i}$ | Average $\Delta\mathrm{RMSE}_{\mathrm{indiv}}$ | Std. $\Delta\mathrm{RMSE}_{\mathrm{indiv}}$ | Minimum $\Delta\mathrm{RMSE}_{\mathrm{indiv}}$ | Maximum $\Delta\mathrm{RMSE}_{\mathrm{indiv}}$ |
|---|---|---|---|---|---|---|---|
| M41 | 100 | Inferior | 0.025 | 0.033 | 0.002 | 0.030 | 0.037 |
| M41 | 100 | Central | 0.025 | 0.035 | 0.002 | 0.032 | 0.039 |
| M41 | 100 | Superior | 0.034 | 0.044 | 0.002 | 0.041 | 0.048 |
| M41 | 75 | Central | 0.024 | 0.031 | 0.002 | 0.026 | 0.034 |
| M79 | 75 | Inferior | 0.028 | 0.029 | 0.002 | 0.025 | 0.032 |
| M79 | 75 | Central | 0.033 | 0.028 | 0.002 | 0.025 | 0.032 |
| M79 | 75 | Superior | 0.020 | 0.025 | 0.003 | 0.020 | 0.029 |

**Table S1**

Captions:

**Figure S1**

**Condition Number Comparison of Biexponential and Monoexponential Models.**

To illustrate the stability of the biexponential model compared to the monoexponential model, we calculated the condition number (CN) of the Jacobian, $J$, for the biexponential and monoexponential models. $J$ was calculated as described in Section 2.1.4 and then evaluated by "numpy.linalg.cond()" function to determine the CN. For each parameter combination, $J$ is unique. Here, we show how fixing all parameters except one of the transverse decay constants changes the CN. With $T_{22}$ free to vary, the other biexponential parameters were fixed at 0.3, 0.7, and 60. The coefficient $c$ of the monoexponential model was set at 0.7. As seen, the CN for the monoexponential is less than that for the biexponential as if the first exponential of the biexponential was removed. It is important to note that the monoexponential CN is consistently less than the biexponential CN. In addition, when $T_{22}$ approaches $T_{21}$ for the biexponential, the CN rapidly increases indicating the extreme ill-posedness of parameter estimation in this regime.

**Figure S13**

**Sample Improvement in MWF Estimation, $\Delta\text{RMSE}_{indiv}$, for Three Individual Noise Realizations of m41 with SNR = 100.**

Panel **(A):** $\Delta\text{RMSE}_{\text{indiv}}$ = 0.034, an 18% improvement in MSE. Panel **(B):** $\Delta\text{RMSE}_{\text{indiv}}$ = 0.036, an 19% improvement in MSE. Panel **(C):** $\Delta\text{RMSE}_{\text{indiv}}$ = 0.033, a 17% improvement in MSE. The positive values and the dominant blue coloration of the regularized pixels in each panel indicate the improvement in parameter estimation with regularization for a single noise realization.

**Figure S14**

**Effect of Regularization on Standard Deviation in MWF Estimation.**

Panel **(A)**: $\Delta\text{STD}_{\text{all},i}$ across 20 noise realizations for a central slice of m41 with a target SNR of 100; the mean value across regularized pixels is 0.034. Panel **(B)**: $\Delta\text{STD}_{\text{all},i}$ across 20 noise realizations for a middle slice of m41 with a target SNR of 75; the mean value across regularized pixels is 0.022. Panel **(C)**: $\Delta\text{STD}_{\text{all},i}$ across 20 noise realizations for a middle slice of m79 with a target SNR of 75; the mean value across regularized pixels is 0.020. The consistent blue pixels and positive $\Delta\text{STD}_{\text{all},i}$ average value indicate decreased variance with regularization through use of $\lambda$ -NL-RR.

**Figure S15**

**Effect of Regularization on Absolute Bias in MWF Estimation.**

Panel **(A):** $\Delta|\text{BIAS}|_{\text{all},i}$ across 20 noise realizations for a middle slice of m41 with a target SNR of 100; the mean value across regularized pixels is 0.003. Panel **(B)**: $\Delta|\text{BIAS}|_{\text{all},i}$ across 20 noise realizations for a middle slice of m41 with a target SNR of 75; the mean value across regularized pixels is 0.014. Panel **(C)**: $\Delta|\text{BIAS}|_{\text{all},i}$ across 20 noise realizations for a middle slice of m79 with a target SNR of 75; the mean value across regularized pixels is 0.020. The mostly positive values for $\Delta|\text{BIAS}|_{\text{all},i}$ indicate that application of $\lambda$ -NL-RR often reduces bias.

**Figure S16**

**Voxel-wise Improvement in MWF Estimation for m41.**

This figure is analogous to Figure 6 but is applied to the axial slices of m41 for a target SNR of 100. Panel **(A)**: an inferior axial slice with an average $\Delta\mathrm{RMSE}_{\mathrm{all},i}$ of 0.025. Panel **(B)**: a central axial slice with an average $\Delta\mathrm{RMSE}_{\mathrm{all},i}$ of 0.025. Panel **(C)**: a superior axial slice with an average $\Delta\mathrm{RMSE}_{\mathrm{all},i}$ of 0.034. Similar to Figure 6, the dominant blue coloration indicates that regularization consistently reduces the RMSE.

**Figure S17**

**Voxel-wise Improvement in MWF Estimation for m79.**

This figure is analogous to Figure 6 but is applied to the axial slices of m79 for a target SNR of 75. Panel **(A)**: an inferior axial slice with an average $\Delta\mathrm{RMSE}_{\mathrm{all},i}$ of 0.028. Panel **(B)**: a central axial slice with an average $\Delta\mathrm{RMSE}_{\mathrm{all},i}$ of 0.033. Panel **(C)**: a superior axial slice with an average $\Delta\mathrm{RMSE}_{\mathrm{all},i}$ of 0.020. As in Figures 6 and S16, the dominant blue coloration indicates that regularization consistently reduces the RMSE.

**Figure S18**

**Regularization Parameter Impact on Variance.**

Panel **(A)**: variance of $c_1$. Panel **(B)**: variance of $c_2$. Panel **(C)**: variance of $T_{21}$. Panel **(D)**: variance of $T_{22}$. To supplement the comparison of MSE values with CRLB measures seen in Figure 2, we present the variance associated with these results. This simplifies the comparison with CRLB and bCRLB measures, although MSE remains the metric of choice. We also plot bCRLB as a function of $\lambda$, bCRLB($\lambda$), providing a comparison reference for var($\lambda$). The CRLB and bCRLB measures are unchanged from Figure 2. As expected, the experimental variance, var($\lambda$), goes to 0 as $\lambda$ becomes large, as is also seen for bCRLB. Consistent with its definition, bCRLB($\lambda$) generally lies below var($\lambda$); we attribute the small region where this relationship is violated to non-convergence of the bCRLB($\lambda$) calculation, as also seen in panel **(D)** of Figure 3. Additional parameter relationships are as previously discussed.

**Table S1**

**Summary Metrics of MWF Improvement.**

The differences in RMSE due to regularization are collected in this table. For every SNR, slice, and participant combination, the values are positive, indicating that regularization consistently reduces the MSE in estimating MWF. The average $\Delta\mathrm{RMSE}_{\mathrm{all},i}$ is calculated as the average of voxels where each voxel has a $\Delta\mathrm{RMSE}_{\mathrm{all},i}$ value. The average $\Delta\mathrm{RMSE}_{\mathrm{indiv}}$ is calculated as the average of 20 noise realizations where each noise realization has a $\Delta RMSE_{indiv}$ value. The low standard deviation of $\Delta RMSE_{indiv}$ values indicate stability of the regularized analysis across noise realizations.
